# 3D viscoelastic drag forces contribute to cell shape changes during organogenesis in the zebrafish embryo

**DOI:** 10.1101/2021.02.23.432503

**Authors:** Paula C. Sanematsu, Gonca Erdemci-Tandogan, Himani Patel, Emma M. Retzlaff, Jeffrey D. Amack, M. Lisa Manning

**Author notes:** Corresponding author, Department of Physics, Physics Bldg. 229B, Syracuse University, Syracuse, New York 13244.

## Abstract

The left-right organizer in zebrafish embryos, Kupffer’s Vesicle (KV), is a simple organ that undergoes programmed asymmetric cell shape changes that are necessary to establish the left-right axis of the embryo. We use simulations and experiments to investigate whether 3D mechanical drag forces generated by the posteriorly-directed motion of the KV through the tailbud tissue are sufficient to drive such shape changes. We develop a fully 3D vertex-like (Voronoi) model for the tissue architecture, and demonstrate that the tissue can generate drag forces and drive cell shape changes. Furthermore, we find that tailbud tissue presents a shear-thinning, viscoelastic behavior consistent with those observed in published experiments. We then perform live imaging experiments and particle image velocimetry analysis to quantify the precise tissue velocity gradients around KV as a function of developmental time. We observe robust velocity gradients around the KV, indicating that mechanical drag forces must be exerted on the KV by the tailbud tissue. We demonstrate that experimentally observed velocity fields are consistent with the viscoelastic response seen in simulations. This work also suggests that 3D viscoelastic drag forces could be a generic mechanism for cell shape change in other biological processes.

**Highlights:**

- new physics-based simulation method allows study of dynamic tissue structures in 3D
- movement of an organ through tissue generates viscoelastic drag forces on the organ
- these drag forces can generate precisely the cell shape changes seen in experiment
- PIV analysis of experimental data matches simulations and probes tissue mechanics

**Graphical abstract:** 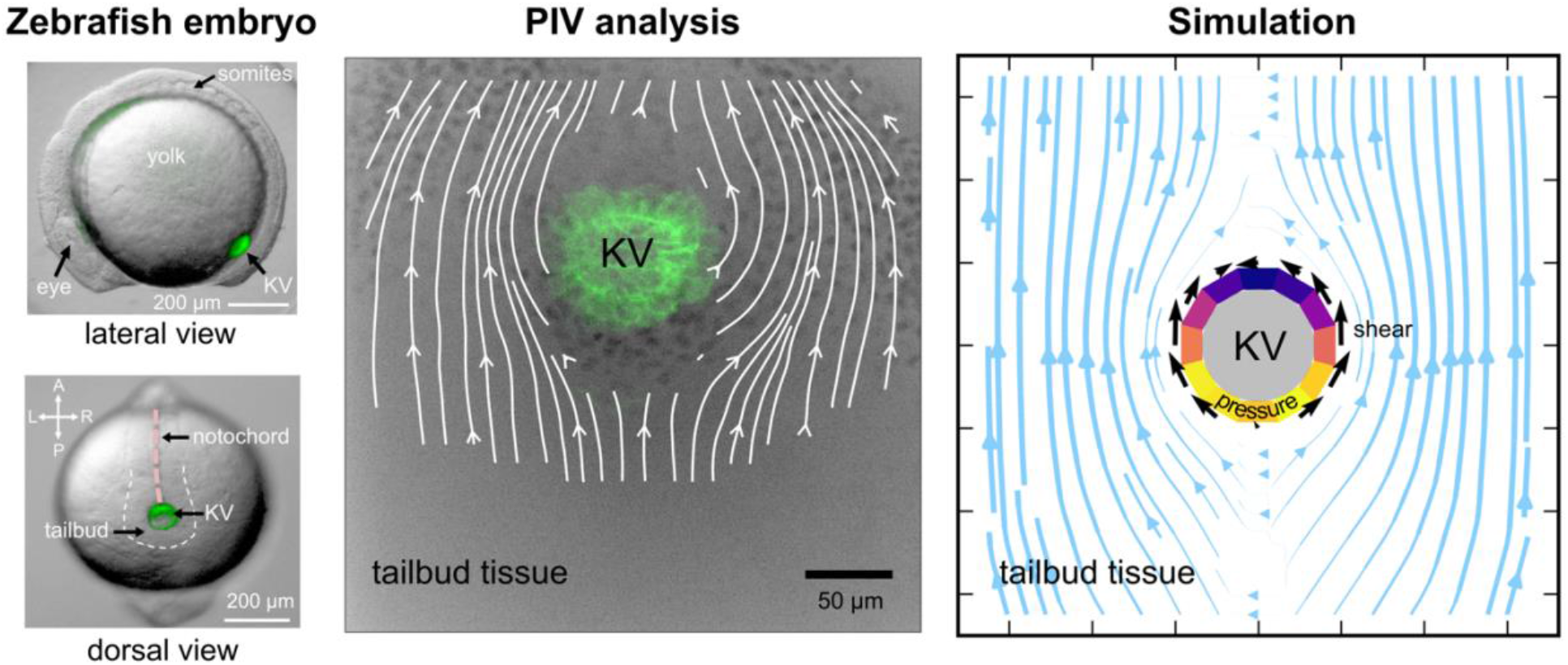

## 1. Introduction

During the early stages of vertebrate embryonic development, the correct establishment of the left-right (LR) body axis is crucial for patterning, positioning, and morphogenesis of several internal organs. When this process is defective, congenital diseases can occur [1] and, in extreme cases, become fatal to the embryo.

Many vertebrate embryos develop a specialized ciliated organ that organizes LR patterning, such as the ventral node in mice, the gastrocoel roof plate in frogs, and Kupffer’s vesicle (KV) in zebrafish [2]. Therefore, an important open problem is identifying the mechanisms that drive the correct formation of these simple organs, and describing how such mechanisms break down in congenital disease.

Here, we focus on the KV, the zebrafish LR organizer [3, 4], as zebrafish are an excellent model organism: they reproduce rapidly and externally, are optically transparent for microscopy, and are amenable to many genetic and mechanical perturbations. The KV is a transient spheroid, rosette-like organ, located in the tailbud of the zebrafish embryo (Figure 1A,B), and is composed of ~50 monociliated epithelial cells that enclose a fluid-filled lumen [3, 5]. The nascent lumen is visible at 10.5 h postfertilization (hpf) at the 2 somite stage (ss) when dorsal-ventral (DV) and anterior-posterior (AP) axes are already established. As the KV travels through the tailbud tissue from 2 ss to 8 ss (a period of 3 h), the lumen expands, cilia elongate, and KV cells undergo asymmetric cell shape changes along the AP axis. This is best observed in the middle plane of the organ where cells in the anterior region become thin and elongated, and posterior KV cells become flattened and wide [6, 7] (Figure 1C).

**Figure 1:**
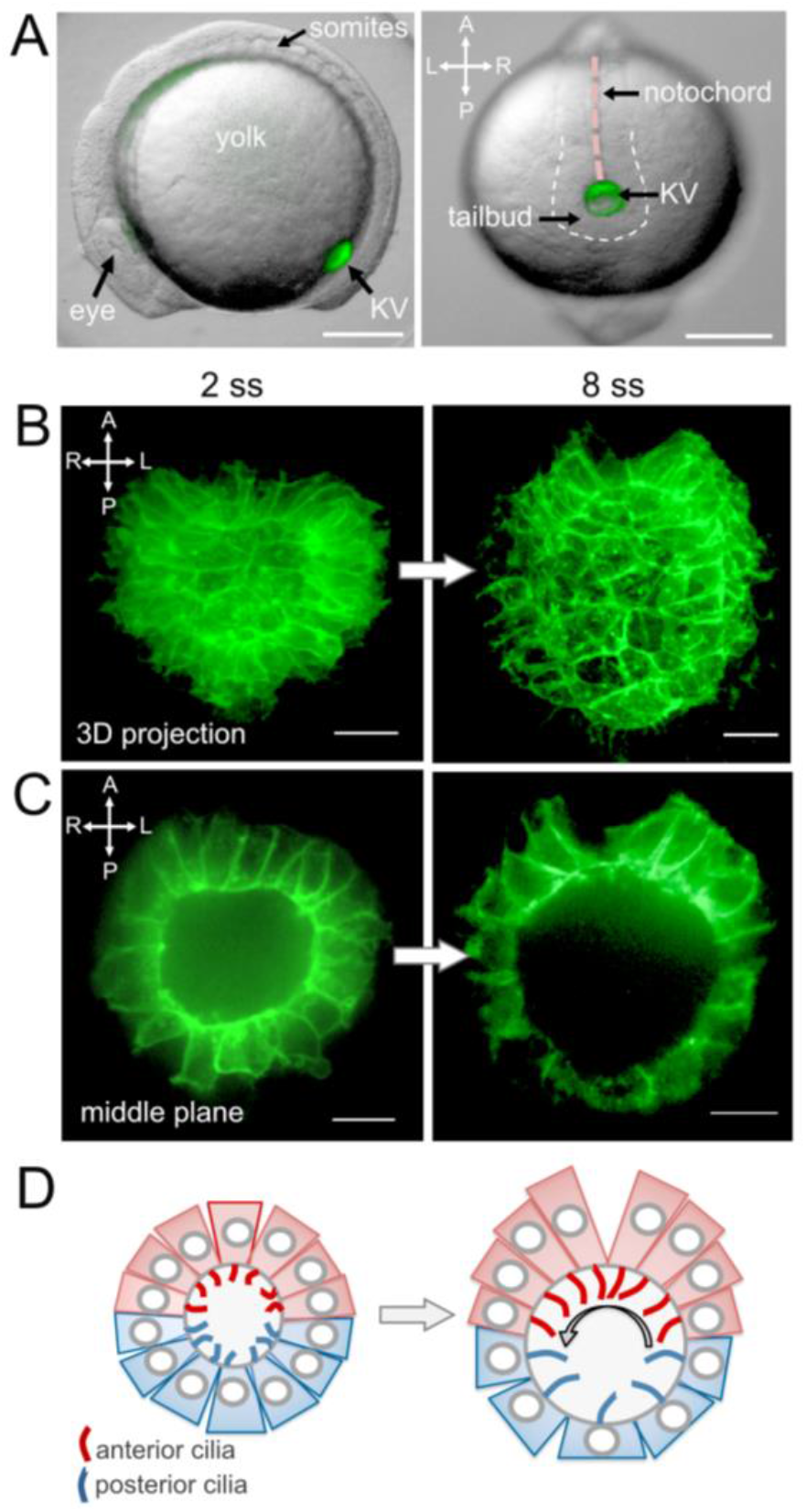
Zebrafish KV cell shape changes during organogenesis between 2 and 8 ss. (A) KV (green) in the zebrafish embryo is located at the posterior end of the notochord in the tailbud seen in the (left) lateral view and (right) dorsal view (scalebar = 200 *μm*). Dotted lines outline the approximate boundary of the tailbud. (B,C) Confocal microscope images (scalebar = 20 *μm*) of the KV dorsal side at (left) 2 ss with symmetric cells shapes and (right) 8 ss with asymmetric cell shapes along the AP axis (note: because the confocal microscope captures mirror images, the left-right axis is flipped). (B) 3D projection of the KV. (C) Middle plane projection. (D) Schematic of KV middle plane cells and cilia (left) at 2 ss when they are AP symmetric and (right) at 8 ss with more ciliated cells on the anterior side that create directional fluid flow to establish LR asymmetry. A=anterior, P=posterior, L=left, R=right.

These programmed changes to cell shape have been shown to be necessary for proper KV function. Specifically, they ensure that there are more ciliated cells packed into the anterior hemisphere compared to the posterior [7]. This asymmetric distribution of monociliated cells along with coordinated KV cilia tilting then generates a directional right-to-left flow across the KV anterior region that develops between 4 to 6 ss and results in stronger forces on the left side and weaker forces on the right side [7, 8] (Figure 1D, right). It has been proposed that stronger mechanical forces on the left side of the KV are sensed by immotile cilia and/or other mechanosensory mechanisms that then biases expression of the *southpaw* gene, which encodes a Nodal signaling molecule, in the left lateral plate mesoderm by 10-12 ss. This asymmetric Nodal signaling is thought to guide subsequent visceral organ morphogenesis [3, 4, 9]. Previous work indicates that the higher density of cilia on the anterior region of the KV is necessary to generate sufficient flow strength for the downstream LR readout [8, 10].

Although programmed shape changes in the KV have been identified as an upstream driver of directional fluid flow and subsequent LR patterning [7, 8, 11], the precise mechanisms that drive KV cell shape changes remain unclear. Previous work has demonstrated cell shape changes depend on an asymmetric accumulation of extracellular matrix at the anterior pole [12], and occur independent of lumen expansion [6]. Additional work indicates shape changes require non-muscle myosin II activity, which regulates actomyosin-based contractility [11]. However, the signals that drive differential shape changes remain unknown. Despite the lack of evidence for specific pathway(s) controlling KV cell shape changes, it is not our intent to rule out the involvement of signaling in this process.

Recently, some of us Erdemci-Tandogan, Clark [13] proposed that perhaps a mechanical signal, generated by the KV traveling through tailbud tissue toward the posterior end of the embryo, could impact shape changes. Specifically, a 2D model demonstrated that drag forces from tailbud tissue are a plausible generator of AP asymmetry in the KV. In experiments, we also demonstrated that the average velocity of the KV is higher than the surrounding tailbud tissue, indicating that the KV has a velocity relative to tailbud tissue. However, there was no direct evidence of drag forces from experimental data, as that work did not quantify the velocity gradients that are responsible for drag.

Moreover, that work focused entirely on a 2D model, but drag forces in 2D are inherently different than 3D at low Reynolds number (*Re* << 1) [14]. This corresponds to flow regimes where viscous forces dominate over inertial forces, also called Stokes or creeping flow. Therefore, a 3D model for the system would be valuable, though technical challenges have, for the most part, inhibited the development 3D models, especially in confluent systems with multiple tissue types.

Such a 3D model would allow us, for the first time, to quantitatively test our hypothesis that tissue flow and drag forces around the roughly spherical KV resemble those of Stokes flow around a solid sphere. Figure 2 shows a 2D cross-section of the analytic solution for Stokes flow around a 3D sphere, illustrating the streamlines (lines of constant velocity in the surrounding fluid, blue lines), mechanical shear stresses at the boundary (black arrows), and pressure on the sphere (colored ring). Gradients in the strain rate (which are perpendicular to the blue lines) generate drag forces that result in pressures (pushing inward on the sphere at the leading edge and outward on the sphere at the trailing edge) and shear stresses (maximum on the sides of the sphere – left and right sides in this representation), that could potentially alter the global shape of the sphere as well as individual cell shapes that comprise the sphere.

**Figure 2:**
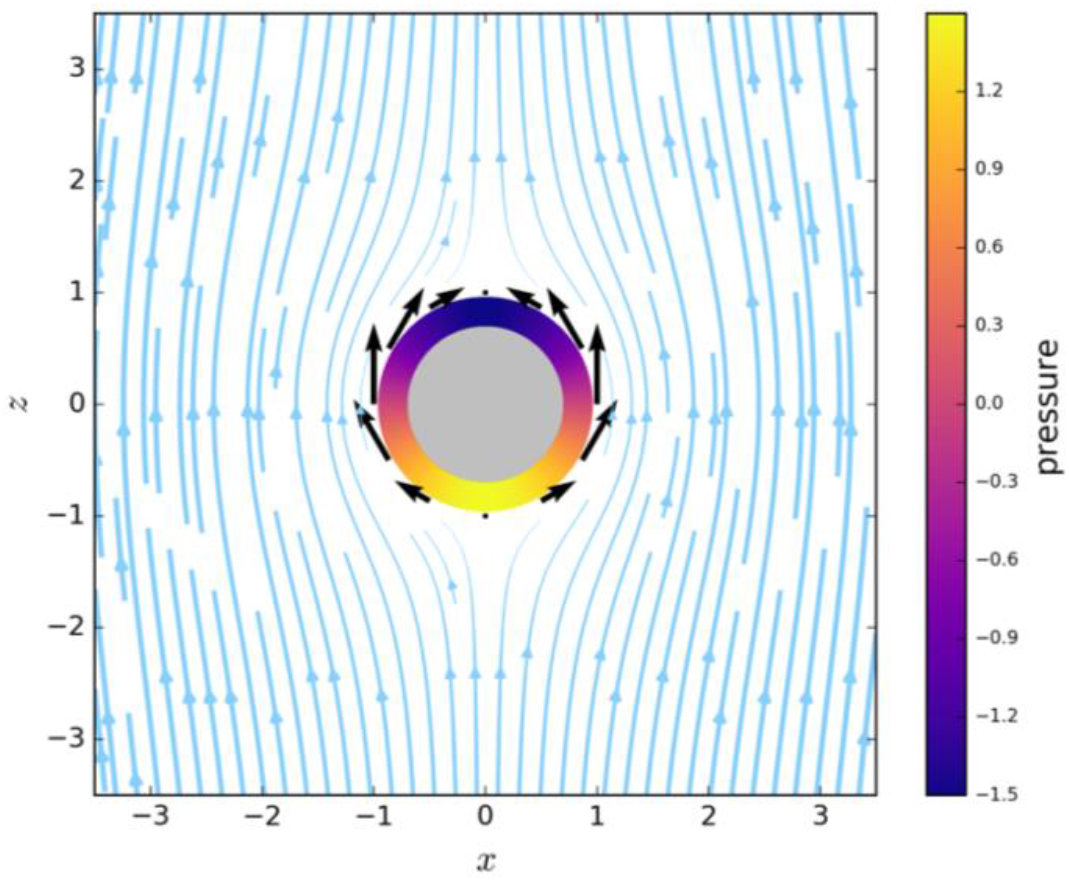
Analytical solution of Stokes flow around a sphere. The colored ring represents the pressure exerted by the fluid on the sphere, with positive pressure (pushing inward) at the bottom of the sphere, upstream of the flow, and negative pressure (pulling outward) at the top of the sphere, downstream of the flow. Arrows represent the shear stress acting tangentially on the KV (the arrow length represents the magnitude, and the location of the shear stress is at the tail of the arrow).

Simultaneously, evidence has emerged that the mechanical response, or rheology, of zebrafish tailbud tissue is quite complex. Therefore, a second open modeling challenge is characterizing the rheology of the tailbud and investigating drag forces in this complex tailbud “material”. Recent experimental work has shed some light on the issue. Mongera, Rowghanian [15] demonstrated that there is a strong gradient in the zebrafish tailbud tissue mechanics, ranging from the more anteriorly located presomitic mesoderm, which is solid-like, to the posteriorly located progenitor zone, which is fluid-like.

To explain the physiological relevance of this transition, Banavar, Carn [16] developed a computational model showing that a fluid-to-solid transition is necessary for proper morphogenesis. They used a coarse-grained continuum approach to model zebrafish body axis elongation with and without a fluid-solid transition of the tissue and demonstrated that unidirectional body elongation occurs only when a fluid-solid transition exists.

While valuable, such large-scale continuum models specify the solid-like or fluid-like locations in a tissue *a priori*, and they are disconnected from experimental observations. In contrast, other cell- and particle-based models have been developed which also exhibit a fluid-solid transition as a function of model parameters, but these model parameters are at the scale of a cell and can be inferred directly from experiments. For example, vertex and Voronoi models for confluent tissues undergo a phase transition as a function of the dimensionless cell surface area parameter *s*_0_, defined as the ratio of cell area to cell volume, 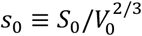 (see details in Methods), exhibiting a solid-like phase when *s*_0_ < 5.413 and liquid-like phase for s_0_ above this value [17]. Particle-based or foam-like models [18, 19] show similar transitions as a function of packing fraction or number density.

Regardless of whether a vertex-based, particle-based, or continuum-based model (such as in [16]) best describes the tailbud tissue, it is important to systematically understand the effect of tailbud tissue rheology on drag forces [20, 21]. Although previous experimental work measured local rheology of the tailbud tissue by generating small perturbations to cell-scale droplets [22], there are no experimental estimates of the effective rheology of the tailbud tissue caused by large-scale perturbations that might be produced, for example, by an organ moving through the tissue. It is well-established that the large-scale effective rheology of a material can be quite different from the rheology measured at small scales [23].

These questions are important beyond just the question of KV in zebrafish development. In many developmental processes, groups of cells must migrate through other tissues to reach a destination and perform a function. If such migratory behavior and corresponding drag forces do generate robust cell shape changes, such shape changes could play an important role in shaping organs in many other contexts. More broadly, such mechanical signaling provides a plausible mechanism by which organisms can transduce molecule-scale information and use it to organize structure on the scale of tissues, and conversely allow tissue-scale mechanics to drive shape changes at the level of single cells, creating a feedback loop across length scales.

In this work, we develop a novel 3D model of confluent tissues to simulate KV morphogenesis, which represent a significant advance from previously studied 2D models. We use these 3D simulations in combination with particle image velocimetry (PIV) analyses of live images of KV development in zebrafish embryos as shown in Supplementary Video S1. We demonstrate that the KV experiences drag forces as it moves through the tailbud tissue and quantify how these mechanical forces contribute to KV cell shape changes. In addition to characterizing velocity fields of tailbud tissue flow around the KV, we use the moving KV as a probe to characterize the effective tissue rheology of the model, for various values of model parameters. Finally, we compare simulation and experimental velocity gradients to place bounds on the model parameter regimes that appropriately capture KV dynamics and mechanics in the zebrafish embryo.

## 2. Methods

### 2.1 Zebrafish embryo imaging

In this study we used the *Tg(sox17:GFP-CAAX)^sny101^* transgenic strain of zebrafish that expresses membrane-localized GFP in KV cells [6]. To simultaneously visualize KV and the nuclei of surrounding tailbud cells, *Tg(sox17:GFP-CAAX*) embryos were injected between the 1 and 2-cell stage with 10 pg of synthetic mRNA that encodes mCherry with a nuclear localization sequence (nls-mCherry). The nls-mCherry mRNA was synthesized using a mMessage mMachine SP6 Polymerase kit (Thermo Fisher Scientific). Injected embryos were incubated at 31°C until early somite stages of development, and then manually dechorionated and mounted in 1% low-melting-point agarose in glass-bottomed dishes (MatTek). Successfully mounted embryos were then imaged starting between 2 and 7 ss using a 20X objective on a Perkin-Elmer UltraVIEW Vox spinning disk confocal microscope (Perkin Elmer) while maintained at 32°C. Under these conditions, we found that embryos developed one new somite pair every 25 min. Recently published work [24] shows that incubating the zebrafish strain TL at 33°C reduces KV lumen size and induces heart laterality defects. We note that incubating the *Tg(sox17:GFP-CAAX*) strain used in this study at 32°C does not alter KV lumen size or heart laterality as shown in Figure S1 (for more details, refer to Supplementary Information).

KV in the tailbud was imaged from the dorsal side, but our confocal microscope captures mirror images such that left-right axis is flipped. Images were captured every 2 or 4 min for different durations as shown in Figure 3, which depended on how long the KV and tailbud cells remained in focus (refer to Supplementary Table S1 for imaging details of each embryo). ImageJ software [25] was used to create maximum projections for each timepoint. We use a *z*-depth of 30 *μm* above and 10 *μm* below the KV midplane for each embryo. We compared various *z*-stack depths and settled on 40 *μm* because it better captures tailbud cells that flow near the KV midplane, where cell shape changes occur, and eliminates irrelevant tailbud cells that are far away from the KV. We set *t* = 0 at 2 ss. All experiments using zebrafish were approved by the Institutional Animal Care and Use Committee at SUNY Upstate Medical University. Embryos were staged as described in Kimmel, Ballard [26].

**Figure 3:**
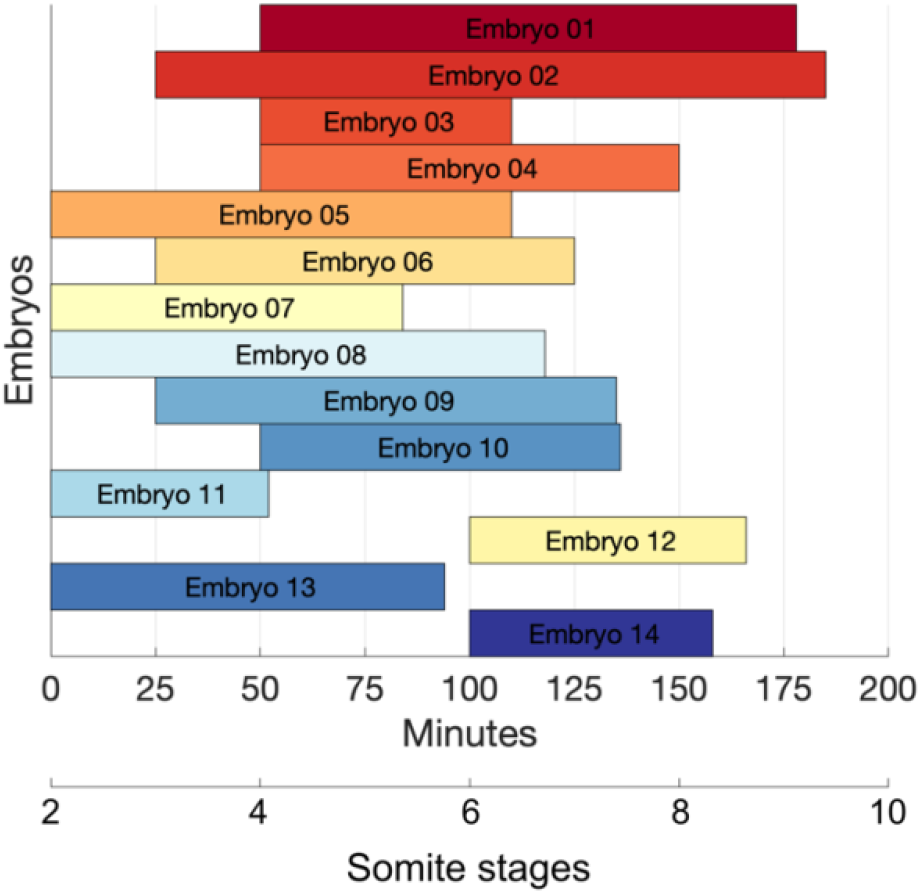
Developmental stage in which embryos images were acquired

### 2.2 Particle image velocimetry (PIV) analysis

To quantify the flows in the tailbud tissue surrounding the KV, we use standard particle image velocimetry (PIV) [27] implemented in the MATLAB PIVlab software package [28]. Additional details are provided in Appendix A.

For the purpose of comparing embryo behavior between different developmental stages, we calculate the average of the AP-directed (or *z*-) component velocity 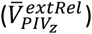 as a function of the distance from the KV along the LR (or *x*-) axis. This velocity profile is computed independently across multiple embryos between 2 to 4 ss, 4 to 6 ss, and 6 to 8 ss, which we will call the *somite average* velocity profile. Besides allowing comparison between developmental stages, the shorter stages spanning 2 ss (rather than 2 to 8 ss) also allow us to reduce the impact of the KV volume expansion on the velocity gradient as the size of the KV plays a role in its surrounding drag forces. Because embryos span different ranges in the *x*-axis, we only average data points that contained at least two embryos. As a way to measure and compare rates of shearing strain (i.e. velocity gradients: 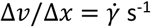; for more details on strain rates, see 3D mathematical model Section 2.3) between various embryos, we translate the data point at the KV surface to the origin and fit a linear profile within error bars using Deming regression [29] implemented by Browaeys [30]. The linear regression only included points within ~50 *μ*m away from the KV surface because we would like to measure rate of shearing strain close to the KV.

### 2.3 3D mathematical model

We use a 3D self-propelled Voronoi (SPV) model [17] to study the movement of the KV through the tailbud tissue. The Voronoi model is a vertex-like model where the degrees of freedom are the cell centers, and cell geometry is defined by a Voronoi tessellation – whereas in vertex models the degrees of freedom are the cell vertices. This main distinction offers two key computational advantages: (a) implementation of cell motility is straightforward in Voronoi models as the self-propulsion is implemented at the cell centers and (b) T1 transitions are intrinsically solved by the Voronoi tessellation. Both models use an energy functional to obtain the preferred geometry of the cells in the system. In 3D SPV model, the energy of a confluent tissue with *N_C_* cells is:

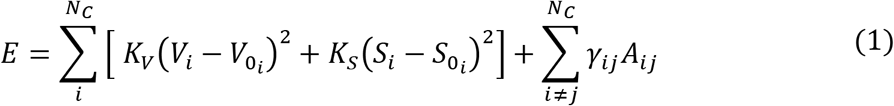

where the first term of the right-hand side (RHS) represents the volume compressibility of a cell with *V_i_* and *V*_0_*i*__. representing actual and preferred volume, respectively. The second term models the actual surface area of a cell, *S_i_*, given the preferred surface area *S*_0_*i*__. This term represents the mechanical forces generated by the competition between cell-cell adhesion molecules and actomyosin in the cytoskeleton localized near the cell cortex. *K_v_* > 0 and *K_S_* > 0 are elastic moduli related to the deviations from the preferred cell volume and surface area, respectively.

Finally, the last term on the RHS, which does not appear in the model reported in [17], represents an additional surface tension, *γ_ij_*, between cells *i* and *j* of different cell types that share a surface area *A_ij_* [31, 32]. Here, we used the heterotypic surface tension *γ_ij_* to create a mechanical boundary between two tissue types. This new term allows us to dynamically simulate coherent tissue structures, such as the KV organ. Compared to the energy functional of [17] that does not include the heterotypic surface tension and the minimum occurs at *s*_0_, the extra tension alters the minimum of the surface tension term. However, it does not abrogate the existence of a minimum and, therefore, we still have a stable system.

The cell dynamics are determined by an over-damped equation with the added self-propulsion velocity, *v*_0_:

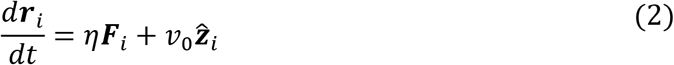

where ***r**_i_* is the position of cell *i, η* is the mobility coefficient (inverse microscopic frictional drag), 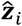 is the direction of the self-propulsion velocity, and ***F*** is the mechanical force on each cell *i*:

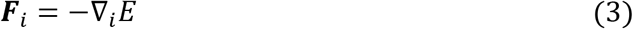

where the energy gradient is given by the energy derivative with respect to a distance vector *r_ij_*:

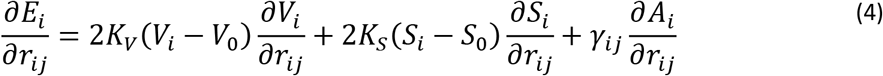

where *∂V_i_/∂r_ij_* and *∂S_i_/∂r_ij_* are the volume and surface derivatives (for details, refer to Appendix A.4.Forces in Merkel and Manning [17])

The self-propulsion velocity *v*_0_ in Equation (2), which is a self-propulsion force times a mobility, is the simplest way to incorporate our observation that the KV moves through the tailbud. It is meant to approximate possible mechanisms that drive this motion. For example, during embryonic development, convergence and extension (CE) movements elongate the notochord [33]. This elongation has been observed in our live imaging and suggests that the notochord “pushes” the KV in the posterior direction through the tailbud cells [13]. Another process that may generate forces on the KV is the active movement of KV posterior cells crawling on tailbud cells and “pulling” the entire KV towards the posterior direction [34]. As we expect that the mechanism that drives KV motion does not strongly affect drag forces that result from such motion, we focus on the simplest self-propulsion model here – investigating other mechanisms would be an interesting avenue for future work.

Our simulations are performed under periodic boundary conditions on a fixed cube with side length *L*. The energy functional is nondimensionalized by the energy unit 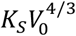, time is nondimensionalized by the natural time unit 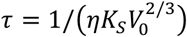, and length is nondimensionalized by 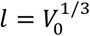. These generate the effective shape index 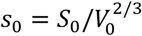. Given these definitions, all simulation results are presented in natural units. As discussed in the introduction, previous work has demonstrated that tissue fluidity is controlled by the shape index. In the limit of low active fluctuations, smaller values of *s*_0_ < 5.413 correspond to solid-like tissues, and increasing *s*_0_ above this value generates fluid-like tissues [17]. Merkel and Manning [17] have shown that this rigidity transition is largely insensitive to *K_V_* values over four orders of magnitude. Although tailbud cells in the zebrafish embryo are self-propelled [34, 35], we are interested in the motion of the KV relative to the tailbud cells. Thus, to simplify the model, tailbud cells are modeled without self-propulsion and we will refer to tailbud tissue as “solid-like” when the shape parameter is lower than that critical value, and “fluid-like” when the shape parameter is higher. However, the motion of the KV through the tailbud could generate interesting dynamic viscoelastic rheological responses in both fluid and solid phases. The exact dynamics remain unknown, as a detailed study of 3D SPV model rheology has never been performed.

In this work, we define three cell types as shown in Figure 4A: external tailbud cells (gray), KV cells (red and blue), and lumen “cells” (yellow). In the zebrafish embryo, the lumen is a fluid-filled vesicle, and in our model, we represent that roughly spherical volume with Voronoi cells (Figure 4A,C). KV is composed of ~50 epithelial cells that surround the lumen [6]. Thus, as illustrated in Figure 4A,B, we used 25 KV anterior and 25 KV posterior cells that are labeled differently solely for post-processing analysis purposes. KV anterior and posterior cells have the same characteristics (i.e., simulation parameters), which means that we do not introduce any innate preferred asymmetry in the simulations. The initial positions of the KV cells are distributed as a spherical monolayer surrounding the lumen. Additional details of the model implementation are provided in Appendix B.

**Figure 4:**
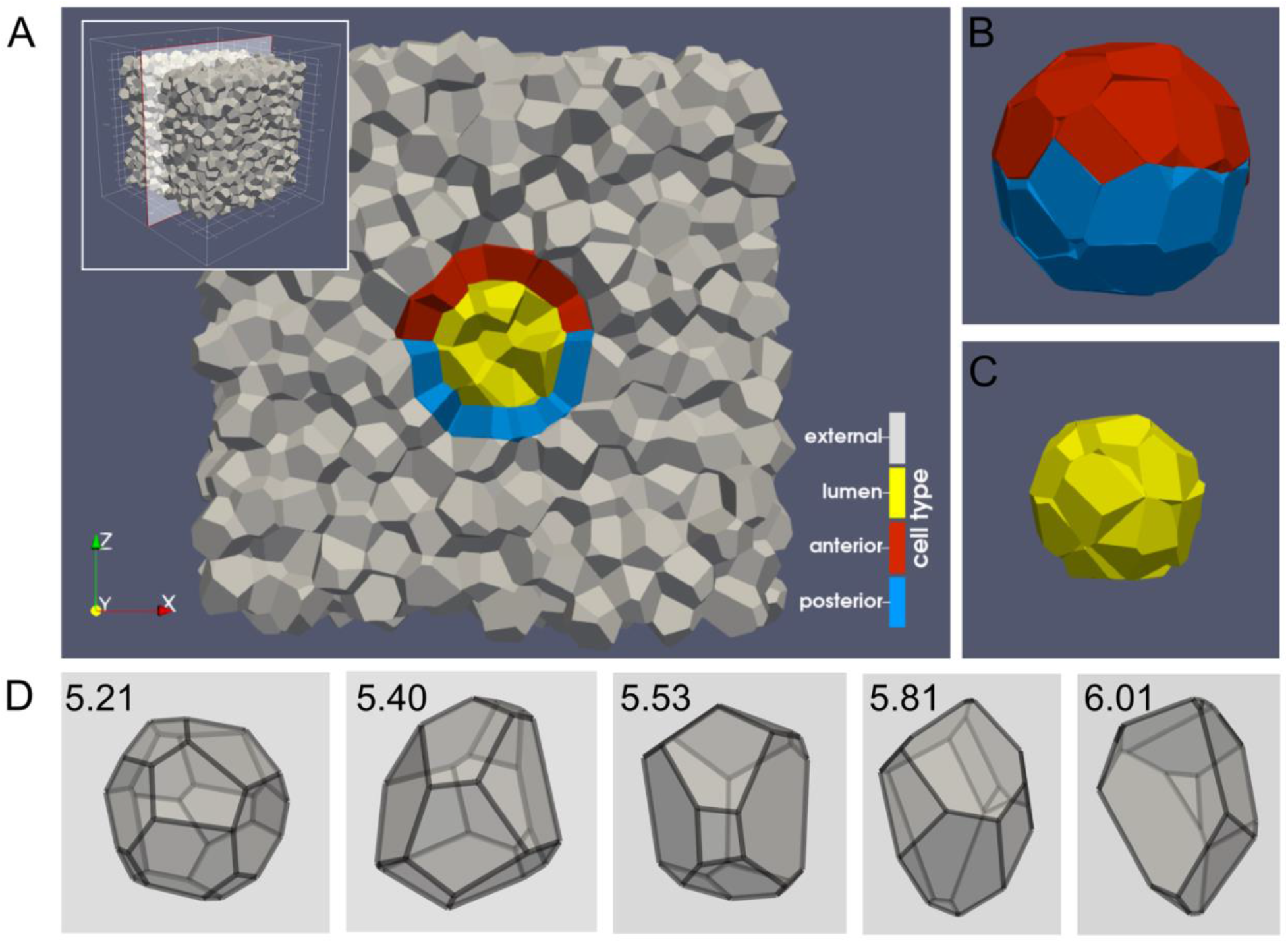
Initial configuration of the 3D SPV model after the system converges to a state of minimal energy and before self-propulsion is introduced. The system contains 2048 external cells, 50 KV cells, and 17 lumen cells. (A) A cutout of the entire computational domain through the center of the KV as shown in the inset. (B,C) Initial spherical shape of the (B) KV and (C) lumen. (D) Increasing values of observed cell shape parameter *s*_0_ with solid-like cells (*s*_0_ < 5.413) more compact and spherical and liquid-like cells (*s*_0_ > 5.413) more elongated.

In previous work by some of us [6, 11, 13], asymmetry between anterior and posterior cells in the KV was quantified using the anterior-posterior parameter (*APA*), which is the difference between the length to width ratio (*LWR*) of anterior and posterior cells (*APA* = *LWR_ant_* – *LWR_post_*) from a 2D cross section. While defining *APA* from a 2D cross section is straightforward, there are several different choices for how to define *APA* when one has access to the full 3D shape [6, 13], and we find that such measurements are generally less robust. Therefore, we turn to a different measurement here. *APA* readout is directly related to the ratio of anterior to posterior cells (*n_a_/n_p_*): when the KV is symmetric at 2 ss, all KV cells have similar shape and each hemisphere has half of KV cells; at 8 ss, anterior cells are thin and elongated whereas posterior cells are wide and flattened, and this results in more cells being located on the anterior than on the posterior side. As the cell distribution readout is more robust in our 3D simulations, and has been shown to drive the lumen asymmetric fluid flows and LR symmetry breaking, we will use *n_a_/n_p_* as the primary readout parameter for KV asymmetry, though we also show data for *APA* in Supplementary Figure S2.

In addition to predicting motion, an additional benefit of physical models is that they can be used to predict forces and stresses. Here, we can use Equation (1) to analyze forces exerted by tailbud tissue on KV cells. Specifically, using Equation (3), we obtain the force ***F**_jj_* of an external cell *i* on a KV cell *j* and then calculate the traction on each KV cell:

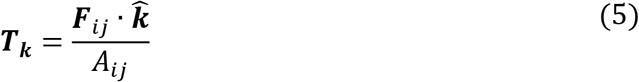

where *A_ij_* is the area of the facet shared between cells *i* and 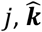 is the unit vector normal 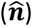 or tangential 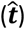 to *A_ij_*. We define 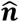 to point out from *A_ij_* and 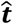 is the vector tangential to *A_ij_* that gives the maximum tangential traction. These tractions can then be used along with velocity gradients to quantify the rheology of the tailbud tissue next to the KV and probe how the drag forces generated by tailbud tissue are stretching and moving cells within the KV. Additional details for how the streamlines and forces are averaged over the unstructured grid in Voronoi models are given in Appendix C.

To better understand the rheology of external cells, we analyze how strain 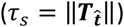 and rate of shearing strain, 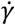, are related for different tailbud tissue fluidity and KV initial velocity. The rate of shearing strain is calculated using the gradient of the two velocity vectors closest to the KV. On the *xz*-plane:

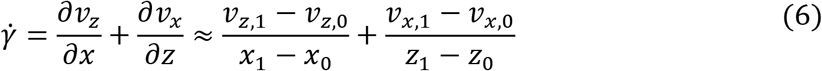

where the index of *v* indicates the velocity components. At the Equator, *v_x_* is negligible and Equation (6) can be simplified to 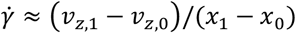.

## 3. Results

### 3.1 Simulations

We use 3D modeling to assess how KV cell distribution varies for a range of values of tailbud tissue fluidity (*S*_0_) and KV initial velocity 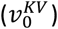. Since we hypothesize that fluid-like drag forces generated by KV motion alters cell shape, we first focus on quantifying the motion of the KV through the tissue as a function of model parameters. Figure 5E illustrates the observed KV steady-state speed 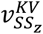 as a function of input KV self-propulsion 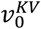 and tailbud tissue mechanics (parameterized by the shape parameter *S*_0_). There is a solid-like regime in the bottom left corner of the phase diagram at low magnitudes of self-propulsion and compact cell shapes, where the KV steady state velocity 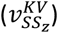 is approximately zero; the self-propulsion is not sufficient to overcome the solid-like tailbud tissue that occur when cell shape indices are small.

**Figure 5:**
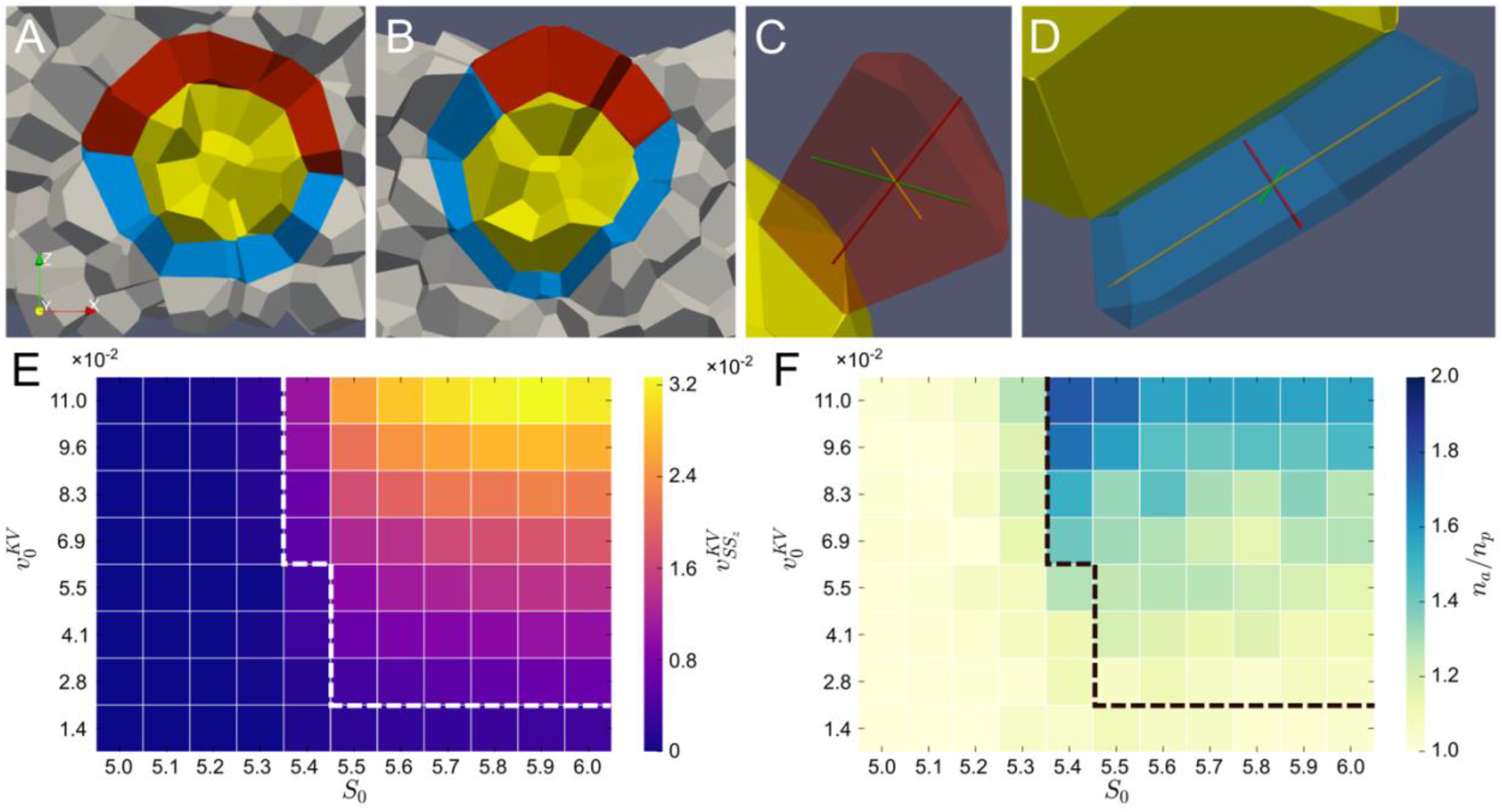
(A,B) Examples of different *n_a_/n_p_* (A) low (*S*_0_ = 5.5 and 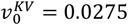), and (B) high *n_a_/n_p_* (*S*_0_ = 5.5 and 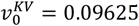). (C,D) Different cell shapes for the KV of panel B: (C) elongated and narrow anterior cell and (D) flattened and wide posterior cell. (E,F) Sweeps in parameter space of tailbud tissue fluidity (*S*_0_) and KV initial velocity (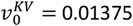, 0.0275, 0.04125, 0.055, 0.06875, 0.0825, 0.09625, and 0.11 labeled for simplicity as 1.4, 2.8, 4.1, 5.5, 6.9, 8.3, 9.6, 11 × 10^-2^): (E) KV steady state velocity 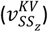 and (F) ratio of number of anterior to posterior cells (*n_a_/n_p_*). The region above and to the right of the dashed lines indicates KV moves in a steady motion and the region below and to the left of the dashed lines indicates either (i) KV does not move or (ii) has very small displacement in solid-like *S*_0_ and 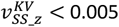.

We also observed very small displacements when 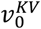 is low and *S*_0_ = 5.4, i.e., when the surrounding tissue is solid-like but close to the fluid-solid transition. In general, this regime resembles an object moving intermittently through a high yield stress solid, instead of through a viscoelastic fluid. In this work, we are interested in the movement of the KV through tailbud tissue as it is observed in the zebrafish embryo: steady movement and with length scales comparable to the KV size. Therefore, we add a threshold to Figure 5E,F (dashed lines) to identify where the KV steady state *z*-velocity 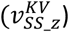 becomes greater than 0.005. While this choice of threshold is somewhat arbitrary, it is sufficiently small to ensure that only regions of model space where 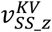 is larger than that threshold are consistent with experimental observations.

In contrast, in the upper right-hand corner of the Figure 5E, corresponding to higher rates of KV self-propulsion and elongated cell shapes that fluidize the tissue, the KV is able to move quickly through the tissue and 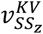 is large. Between these extremes, we observe a smooth gradient in steady state KV speed as a function of model parameters.

Next, we ask how this observed KV motion correlates with anterior-posterior (AP) asymmetric cell shapes changes and distribution of ciliated cells. In contrast to previous results from 2D modeling [13], we observe that the cell shapes defined by the 2D length-to-width ratio (*LWR*) are not as robust in 3D simulations. We hypothesize that perhaps the 2D *APA* = *LWR_ant_* – *LWR_post_* is not good at capturing the full 3D cell shape highlighted in Figure 5C,D (we include data and analyses of the 3D *APA* in Supplementary Figure S2). Since the asymmetric AP distribution of ciliated cells is thought to be the important downstream readout of these shape changes for LR patterning [3], we next investigate whether cells are AP asymmetric, using the ratio of the number of anterior and posterior cells *n_a_/n_p_*. In Figure 5E,F, we see that the distribution of cells in KV remains symmetric (*n_a_/n_p_* ≈ 1) (Figure 5A is an example of symmetric distribution of KV cells) when its steady state speed is approximately zero. This is important, as it confirms that self-propulsion forces alone are not sufficient to generate AP cellular asymmetry; the KV has to move through the tailbud to generate this asymmetry in our model.

In addition, at the slowest self-propulsion speeds, the KV moves so slowly that AP asymmetry is not generated. In the remaining regions of parameter space, the AP asymmetries can become quite large (as exemplified by Figure 5B), and there is a range of parameter values (*S*_0_ ≥ 5.4 and 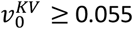) that generate asymmetries similar to published experiments (*n_a_/n_p_* = 1.77) [11] (a simulation in this range is shown in Supplementary Video S2). Interestingly, this panel also highlights an important difference between 2D and 3D simulations: while 2D results [13] showed a monotonic increase of *n_a_/n_p_* as the fluidity of the tailbud increases, the 3D results behave differently, with an enhanced asymmetry when the cell shape index is close to the value for the fluid-solid transition in the limit of low fluctuations (*S*_0_ = 5.4).

Next, we characterize the external cells movement relative to the KV as a fluid flow field and compare it with a known solution of flow around a sphere. In continuous fluid mechanics, a flow is roughly categorized into different regimes according to the Reynolds number, which represents the ratio of inertial to viscous forces: *Re* = *ρUL_Re_/μ*, where *ρ* is the fluid density, *U* is the flow speed, *L_Re_* is a characteristic length scale, and *μ* is the fluid dynamic viscosity. When viscous forces are dominant (i.e. *Re* << 1), by either having flow at very small scales (e.g. microfluidics), very low velocities, and/or very high dynamic viscosity, inertial forces can be neglected, which simplifies the problem to Stokes (or creeping) flow. Given that the KV moves at very slow speeds [13] and its dimensions are in the order of micrometers, we expect that *Re* << 1 in our simulations and compare our simulations to the analytical solution of Stokes flow around a sphere (Figure 2).

To further understand the mechanisms causing the observed enhanced AP asymmetry near the fluid-solid transition (*S*_0_ = 5.4 and 5.5) as well as asymmetries in the region of parameter space where this tissue is ostensibly fluid-like (*S*_0_ = 5.7), we compute streamlines from the velocity field of externals cells and also calculate shear and pressure acting on the KV. First, we compare results of the fluid-like regime with expected streamlines, shear, and pressure of Stokes flow around a sphere (Figure 2). The streamlines in Stokes flow are symmetric both about the *x* and *z* axes, while shear is symmetric about the *z*-axis and its direction is tangent to the streamlines with maximum value at the sphere Equator (*z* = 0). The pressure is also symmetric about the *z*-axis, with positive (pushing inward) upstream pressure (*z* < 0) on the sphere and negative (pulling outward) downstream pressure (*z* > 0).

When the tailbud tissue is fluid-like (*S*_0_ = 5.7) and velocities are intermediate 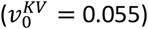, Figure 6F shows the results of SPV model simulations are remarkably similar to that of Stokes flow shown in Figure 2. This demonstrates our hypothesis is plausible for 3D flows: tissue motion surrounding KV can act as a flowing fluid and exert drag forces on the KV via fluid-structure interactions.

**Figure 6:**
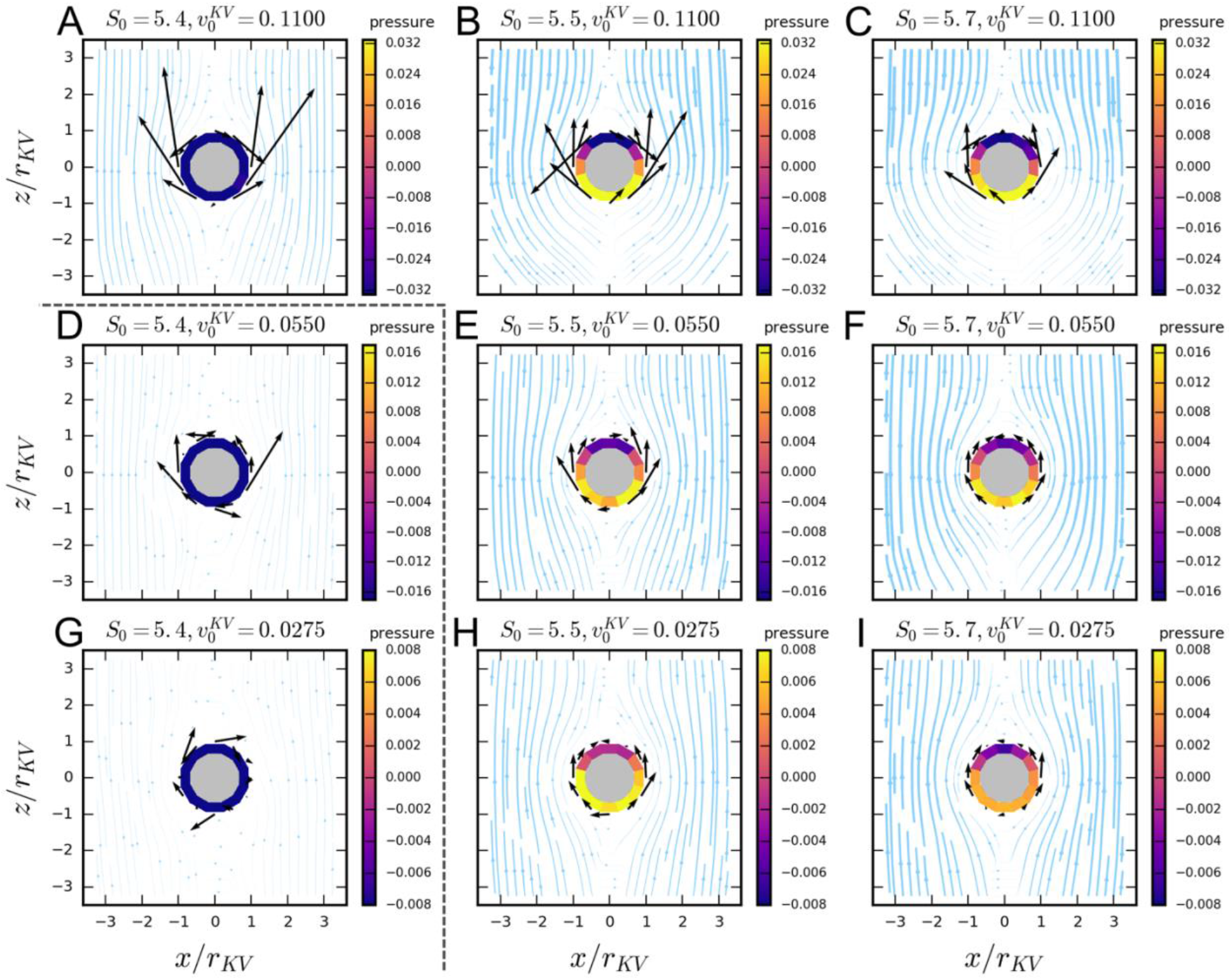
Streamlines, pressure, and shear ensemble average for various tailbud tissue fluidities *S*_0_ and initial KV velocities 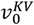. Each row is the result of a fixed 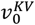 value whereas each column is the result for a given *S*_0_. The colored ring represents the pressure exerted by the external cell fluid flow on the sphere, with positive pressure (pushing inward, yellow-ranged colors) negative pressure (pulling outward, purple-ranged colors). Note that the range of the pressure colorbar is fixed per row and, thus, pressure colors can be compared withing a row. Arrows represent the shear stress acting tangentially on the KV (the arrow length represents the magnitude with the same scale for all plots, and the location of the shear stress is at the tail of the arrow). *x* and *z* axes are normalized the KV radius, *r_KV_* (for more details on this normalization, refer to Appendix C). Streamline width represents the velocity of external cells relative to the KV 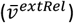 normalized by the initial KV velocity 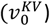 such that streamline widths are comparable within a row. The plots above and to the right of the dashed lines indicates KV moves in a steady motion and the plots below and to the left of the dashed lines indicates KV has very small displacement in solid-like *S*_0_ and 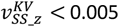.

With increased velocity 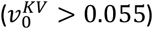 in the fluid-like regime (*S*_0_ = 5.7, shown in Figure 6C), we find that although the pressure was still similar to Stokes’ pressure, streamlines are no longer symmetric about the *x*-axis and shear stresses on the anterior side of KV (*z* > 0) are not tangent to streamlines. The effect of increasing KV velocity on streamline asymmetry can be further seen in Supplementary Figure S3, in which asymmetry about the *x*-axis gradually becomes more pronounced for increasing 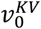 and 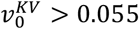. These behaviors that deviate from Stokes flow indicate that the rheology of the tailbud tissue in this parameter regime departs from a Newtonian fluid.

Finally, for weakly solid-like tissues (Figure 6, panels A, D, G) – corresponding to values of the tailbud shape index just below the fluid-solid transition (*S*_0_ = 5.4). Streamlines of Figure 6A and 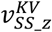 in Figure 5E indicate that external cells do flow around the KV and generate drag forces. However, pressure behavior greatly differs from Stokes flow (Figure 2): pressure is negative both upstream and downstream of the KV. In Figure 6D,G, because the KV is moving very little, small velocity fluctuations due to the local geometry matter a lot and induce quite large fluctuations in the stresses and pressure relative to the mean, resulting in an ensemble average that may be skewed by these large stresses. Thus, we observe shear stress LR asymmetries that are due to these local geometry fluctuations and do not reflect global symmetries. While we include panels D and G for completeness, they are not consistent with experimental observations, and we do not focus on them for the remainder of the manuscript.

A related observation is that for almost all simulation parameters, the velocity of streamlines far away from the KV appears to plateau to a constant value (data points in Figure 7), which is not observed in Stokes flow (thick purple line in Figure 7) in which viscous effects generated by the sphere extend to the far-field external flow. This plateau indicates there are two distinct regions in the external cells surrounding the KV: (1) a yielded region close to the KV and (2) a non-yielded region far from the KV.

**Figure 7:**
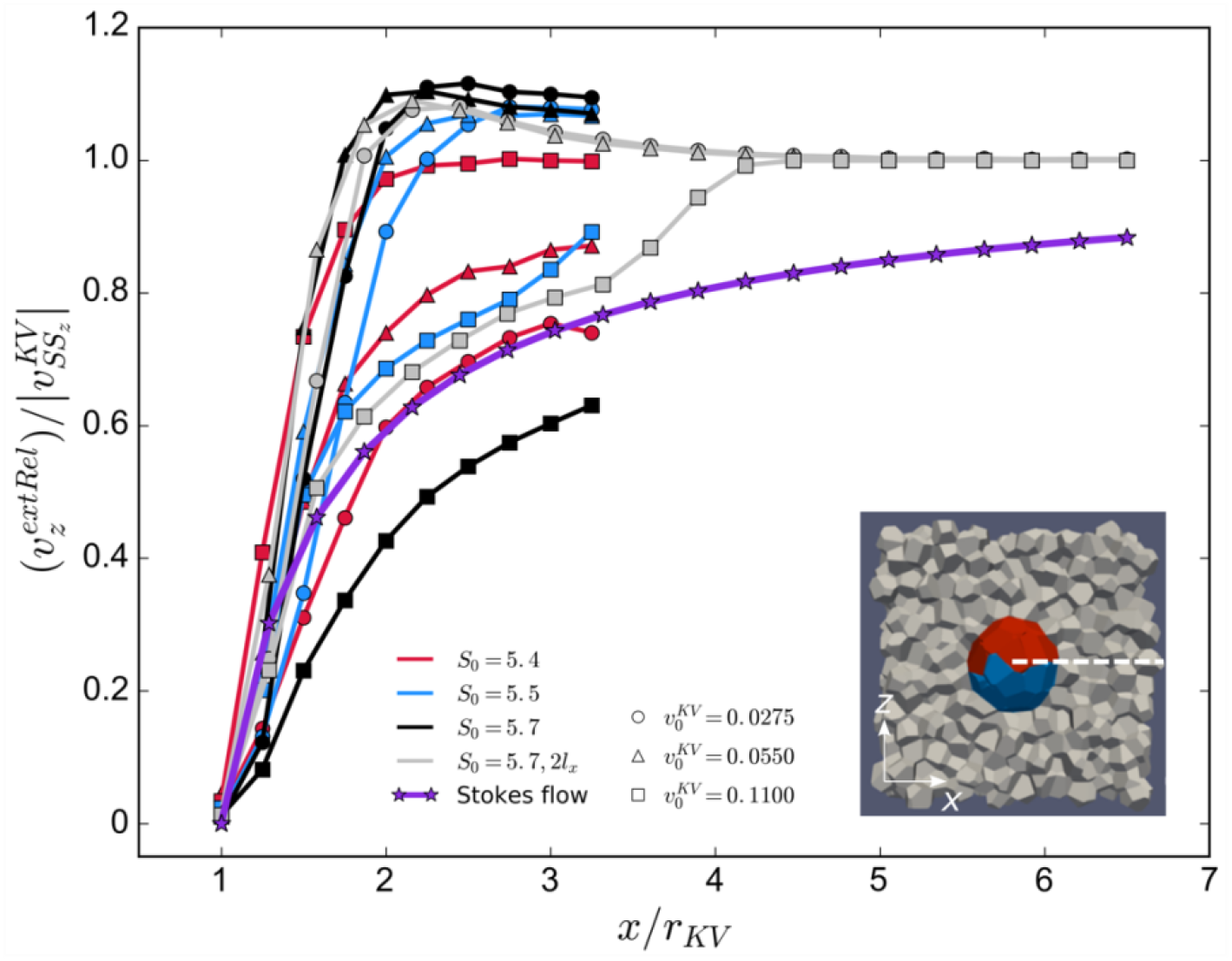
Simulation velocity profiles for different tailbud tissue fluidity (*S*_0_) and KV self-propulsion velocity 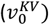. *y*-axis is the external cells *z*-component of velocity relative to the KV, 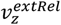, normalized by the KV SS velocity, 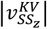. *x*-axis represents the distance from KV center normalized the KV radius, *r_KV_*. Profiles include regular domains of ~3.5*r_KV_* = *l_x_* and larger domain of 2*l_x_* at the Equator as indicated in the inset (see Appendix B for details on *l_x_*).

In summary, we observed the following non-Newtonian behaviors: (a) in the solid-like regime (*S*_0_ = 5.4), KV needs a minimal self-propulsion velocity to overcome forces of external cells otherwise the KV does not move, (b) in the fluid-like regime (*S*_0_ = 5.5 and 5.7) streamline asymmetry about the *x* axis for 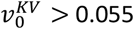, and (c) a non-yielding region in the external cells.

This leads us to hypothesize that in our Voronoi simulations, external cells may behave as a viscoelastic fluid, such as a Bingham or Herschel-Bulkley plastic. In these non-Newtonian materials, a sphere needs a minimal velocity to overcome the materials’ yield stress and start moving [36]. When a sphere moves through Bingham plastic, it creates a yielded (fluidized) region surrounding the sphere and far from it, the fluid remains as a solid [36]. Herschel-Bulkley plastics, in addition to having a minimum yield stress, are also shear thinning so that the material appear less viscous as the rate of deformation increases [37].

We investigate whether external tissues in the simulations have the effective rheology of a viscoelastic material, by analyzing the profiles of external cells velocity relative to the KV 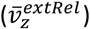 at the KV Equator (*z* = 0) and along the *x* axis (Figure 7). In order to compare our simulation results with results from flow around a sphere, we normalize the simulation velocity 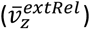 by the absolute value of the KV steady state velocity 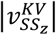, which corresponds to the standard normalization in continuum fluid dynamics. This allows us to study the *shape* of strain gradients, rather than the *magnitude*, as the shape can help us identify the most suitable viscoelastic model for the material rheology.

In the analytic solution to Stokes flow around a sphere, the velocity increases monotonically as a function of distance from the sphere and approaches unity at large distances 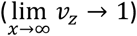. In contrast, in experiments of creeping flow of Bingham plastic around a sphere [38], velocity profiles are characterized by (a) a steeper velocity gradient than Stokes flow, often with an overshoot above unity; and (b) a plateau in the solid region corresponding to unyielded material.

For the velocity profiles corresponding to *S*_0_ = 5.7 in Figure 7, we observe characteristics very similar to that of Bingham plastics, and clearly distinct from a Newtonian fluid. In addition, tissues that would be solid-like in the absence of driving forces (*S*_0_ = 5.4, 5.5) also exhibit Bingham plastic-like features when the KV moves through it, although there is no minimal velocity overshoot.

For computational efficiency most of our simulations are performed for the smallest system size that does not affect the steady-state KV velocity (*N_ext_* = 2048). Figure 7, however, demonstrates that for fluid-like tissues, the velocity profile does not yet attain its plateau value at the edge of the simulation box. Therefore, for *S*_0_ = 5.7, we also simulated larger system sizes (2*l_x_*) to demonstrate that the system does in fact attain a plateau. We note that the velocity profiles for different system sizes do not match because there is a shift in normalization for the smallest system size.

In addition to the velocity profiles at the equator, the velocity profiles of 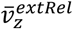 at the two poles and along the *z*-axis, called wake profiles, are frequently used to investigate non-Newtonian behavior. In Stokes flow, the upstream and downstream wakes are symmetric, whereas in viscoelastic regimes the wakes are not symmetric with the downstream wake trails extending over ten sphere radii away from the sphere [39]. We observed a trend of this asymmetry (Supplementary Figure S5), but we do not completely capture the trailing wake due to the simulation domain size of ~3.5 *r_KV_* away from the KV center in the *z*-direction.

In addition to flow patterns, we take advantage of the fact that Voronoi models predict physical forces, and analyze the shear stress *τ_s_* acting at the interface between the KV and external cells, as well as the rate of shearing strain *γ* (i.e. velocity gradients) at the KV-external cell interface, for different model parameters. In Newtonian fluids, the shear stress and the rate of shearing strain have a linear relationship, and the ratio gives the dynamic viscosity of the fluid: 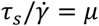. Non-Newtonian fluids, on the other hand, exhibit a non-linear relation between *τ_s_* and 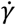, as illustrated in Figure 8A.

**Figure 8:**
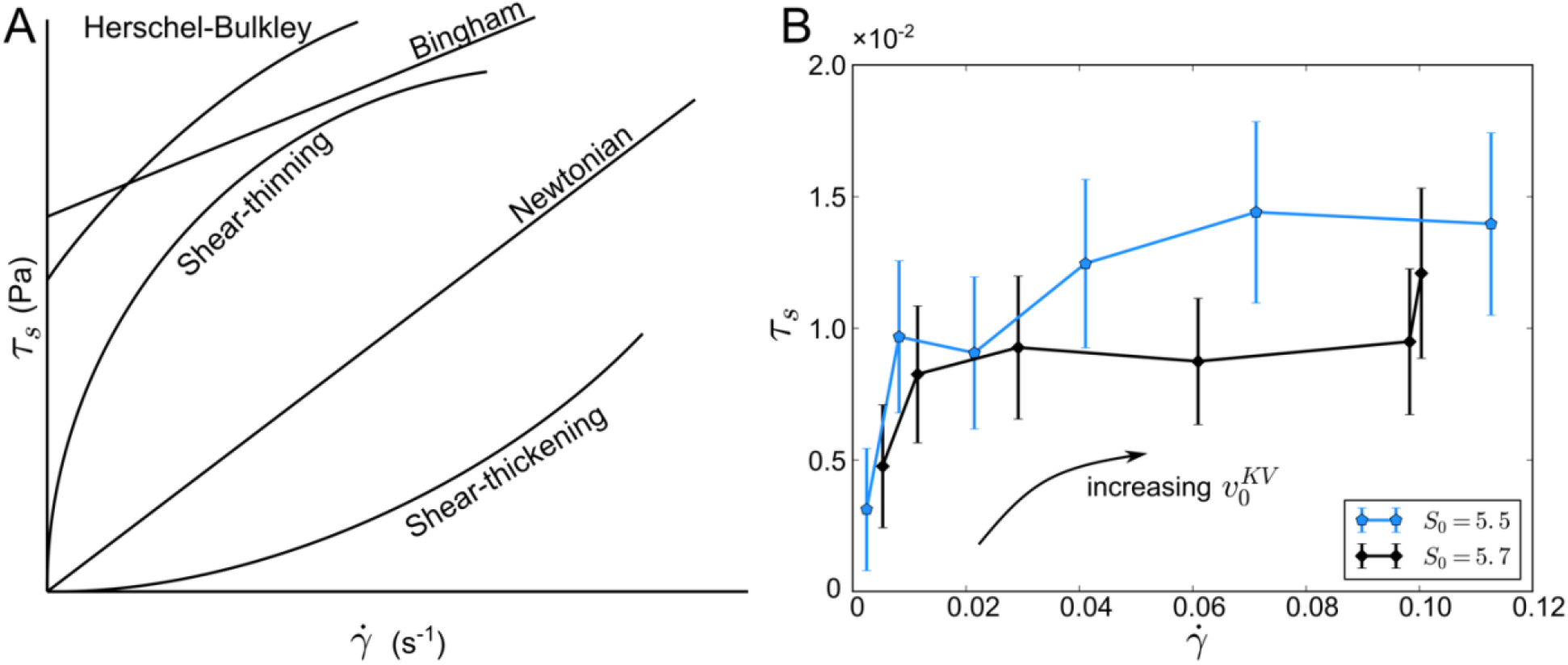
Shear stress (*τ_s_*) as a function of rate of shearing strain (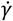, velocity gradients). (A) Curves that represent different viscosity models (adapted from Steffe [37]), where Newtonian fluids are characterized with a linear profile. (B) Curves for two fluid-like tailbud tissue fluidity, *S*_0_ = 5.5 (blue) and 5.7 (black), and increasing 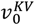 (two largest value excluded) indicate a non-Newtonian behavior, possibly shear-thinning.

Figure 8B presents the relationship between shear stress and strain rate at the equator of the interface of KV and external cells. We see that shear stress generally increases with strain rate, with a non-linear relationship that is clearly distinct from Newtonian flow, and indicates a shear thinning behavior. Although our simulation data is similar to the behavior of epithelial cells monolayer moving around a circular cylinder [40], it is not sufficient to distinguish between various shear-thinning models (e.g. Herschel-Bulkley, Bingham), and this is reserved for future work.

### 3.2 Zebrafish embryos particle image velocimetry (PIV) analysis

To analyze the collective behavior of tailbud tissue *in vivo* and determine whether this tissue acts as fluid flow around the KV such that drag forces are exerted on the KV during morphogenesis, we imaged live zebrafish embryos between 2 and 8 ss (Supplementary Video S1) when KV cells change shape from symmetric to asymmetric along the AP axis [6, 7]. We used *Tg(sox17:GFP-CAAX)* embryos that express green by fluorescent proteins in KV cells and labeled external cell nuclei by expressing nuclear-localized mCherry proteins. We use PIV [27] to quantify the velocity field of the tailbud tissue and KV cells (***v**_PIV_*). This movement, however, is relative to the lab frame and we need to quantify the tailbud (or external) cells movement relative to the KV. Thus, we subtract the KV average velocity from the velocity field 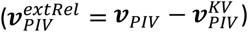 and fix the KV as the reference frame. In Supplementary Video S3, where each frame has been translated in space so that the KV position remains fixed as a reference object, it becomes clear that the tailbud tissue moves around the KV similarly to flow around a sphere.

In Figure 9A, we present the averaged velocity field 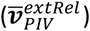 obtained by PIV analysis between 4 and 6 ss of Embryo 10 (Supplementary Video S4) – for an estimation of the PIV analysis error, refer to Supplementary Information and Figure S4. Streamlines of this velocity field (Figure 9B) are remarkably similar to flow around a sphere indicating that drag forces of tailbud tissue are exerted on the KV, which is especially remarkable considering this is tissue moving during embryonic development rather than a well-controlled fluid dynamics experiment. Furthermore, the velocity profile 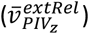 along the LR axis and KV equator (Figure 9C) shows increasing velocity of tailbud cells within ~50 *μ*m away from the KV surface, a clear velocity gradient, after which the velocity stabilizes similarly to the behavior seen in viscoelastic flows around a sphere.

**Figure 9:**
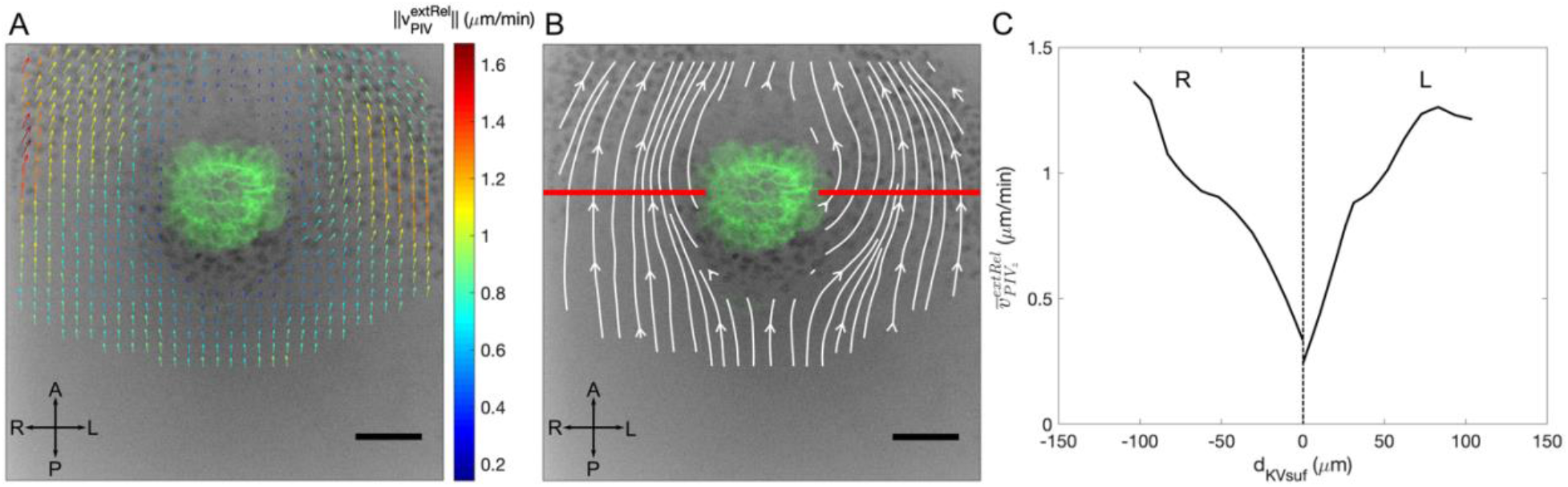
PIV analysis of zebrafish Embryo 10 from 4 to 6 ss with scalebar corresponding to 50 *μ*m. (A) Averaged velocity field. (B) Streamlines that correspond to the velocity field of panel A. (C) Velocity profile of 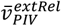 in microns/min along the left-right axis that would pass through the KV centroid as indicated by the red line in panel B.

Due to large variations in biological data, we image 14 zebrafish embryos and evaluate their velocity profiles (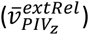 averaged over 2-ss intervals (Figure 10). As expected, embryos’ velocity profiles exhibit variations, though quite a few profiles exhibit the “leveling off” in velocity expected of a viscoelastic material. These variations are completely consistent with embryo-to-embryo variability seen in other features of KV development. For example, published data have shown a high variability between wild-type embryos in KV size and KV cilia number [11, 41]. To test if the gradient variability might be explained by natural variations in the KV, we performed a linear fit on the velocity profiles of individual embryos and compared with results reported by Gokey, Ji [41] (Supplementary Table S2). Given all of the processes happening concurrently such as the embryo’s convergence and extension, the yolk moving along the ventral side of the KV, KV expansion as well as experimental errors (e.g. angle that KV is imaged), the gradients remain remarkably similar within ~25-70% difference depending on developmental stage.

**Figure 10:**
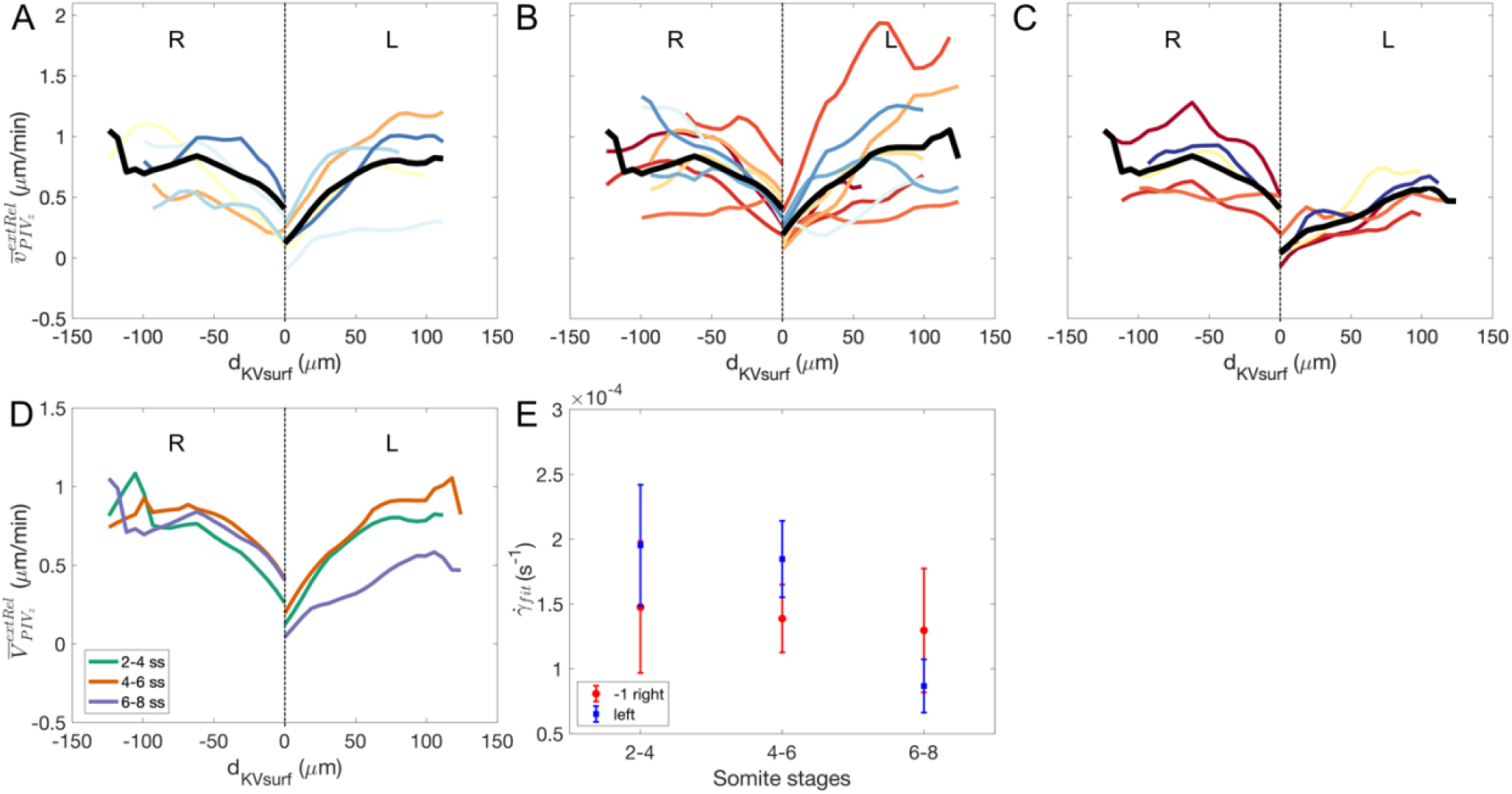
Velocity profiles at the Equator of the KV along the *x*-axis of all embryos over two somite stages. (A-C) Individual embryos (colored lines, where the color represents embryos defined in Figure 3) and the somite average profiles represented by the thick black line. (A) 2-4 ss. (B) 4-6 ss. (C) 6-8 ss. (D) Somite average velocity profiles at each 2 ss interval. (E) Somite average fitted rate of shear strain 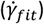, where right profile is multiplied by −1 to facilitate comparison.

Most importantly, we see an undeniable trend of velocity gradients surrounding the KV, which confirms that drag forces are indeed acting on the KV. Moreover, despite relatively large variations in KV size from embryo to embryo[3], the average velocity gradients are remarkably similar across all embryos, and consistent with previously reported results [13]. This robustness lends credence to the idea that the embryos are utilizing these gradients as a mechanical signal, via drag forces.

The overall trend as a function of somite stage can be determined by first quantifying the average velocity profile over all embryos 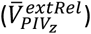) in a given somite stage, illustrated by the black lines in Figure 10A-C, which we term *somite average* velocity profiles and are shown together in Figure 10D to emphasize their differences.

We observe two unexpected but thought-provoking trends in the velocity profiles of Figure 10D: (1) different velocities gradients occur at different somite stages and (2) we see hints of a difference between left and right profiles, with the left-side fluid flow exhibiting a possible shift from stronger velocity gradients (i.e. steeper curves) and generating stronger drag forces at the 2-4 and 4-6 ss to a weaker velocity gradient at the 6-8 ss.

To quantify these observations, we fit a linear profile to the somite averages [29, 30] (Supplementary Figure S6). The slope of the linear fit is the rate of shearing strain 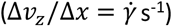 and should be proportional to the drag forces exerted by tailbud tissue on the KV. Because we are interested in the drag forces near the KV, the fit only includes points within ~50 *μ*m from the KV surface. Figure 10E shows the fitted rate of shearing strain 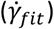 at different somite stages. Both left and right profiles are shown, and negative slopes of the right profiles are multiplied by −1 to facilitate the comparison. We note that the order of magnitude of 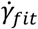 is in agreement with published results that investigate rate of shearing strain between moving cells [42, 43].

In addition, we notice decrease in drag forces only on the left side whereas the right stays fairly constant. This is a striking result considering that, between 2 and 8 ss, it is believed that LR asymmetry has not yet been established. Unfortunately, our data is only suggestive of this new type of LR asymmetry, as velocities profiles are sufficiently noisy that a fluctuation could cause the observed discrepancy. However, this is a very interesting avenue for future research.

Lastly, we would also like to understand whether these velocity profiles provide any insight into the rheology of the tailbud tissue. As discussed earlier in the results Section 3, this can often be determined by studying the shape of the velocity distribution after normalization. In experiments of flow around a sphere, results are typically normalized by the sphere steady-state velocity as done for the velocity profiles of our simulations. For the embryo images, however, the KV moves relative to the tailbud cells as well as relative to the yolk/external reference frame. Thus, PIV analysis gives the KV velocity relative to the external reference frame instead of the tailbud cells. The external cells velocity relative to the KV 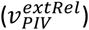 removes the movement of the embryo relative to the external reference frame, and we therefore use the far field value of the velocity profile 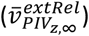 for normalization because the cells furthest away from KV are the least affected by the KV motion – similarly to the far field velocity of Stokes flow or an unyielded region of viscoelastic flows. Additionally, due to the different behavior between the left and right profiles (explained above), we use their respective far field values to normalize the left and right profiles separately.

Figure 11 compares experimental and simulation results by plotting the embryos’ normalized velocity profiles between 4 and 6 ss (thin colored lines) on the same axes as profiles from our numerical simulations of the Voronoi model (thick black and gray lines with markers). See Supplementary Figure S7 for the normalized profiles at 2-4 and 6-8 ss. Natural units of the simulations are converted into microns by using the tailbud cell average diameter ~10 *μm*. The simulation profiles we include here are the ones with the highest (*S*_0_ = 5.7, 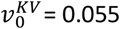) and the lowest velocity gradients (*S*_0_ = 5.7, 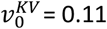) over simulations presented in Figure 7. We also include the larger domain size, 2*l_x_*, (gray lines) which better represents the experimental domain size. This region between velocity gradient extremes encompasses the range of parameter space with enhanced *n_a_/n_p_* asymmetry: the fluid-solid transition (*S*_0_ = 5.4 and 5.5) as well as the region in which the tissue is clearly fluid-like (*S*_0_ = 5.7).

**Figure 11:**
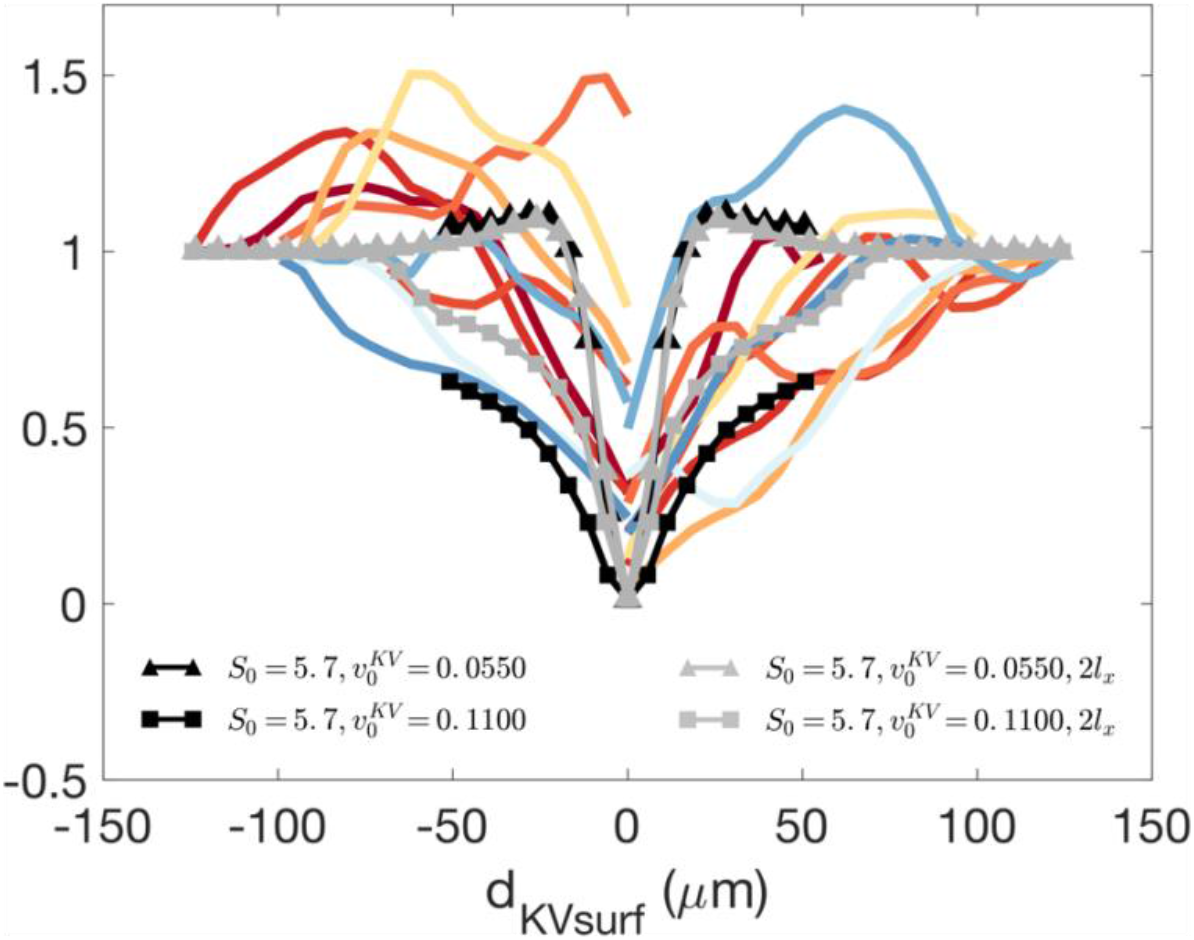
Normalized velocity profiles with embryos represented by colored lines (each color represents embryos defined in Figure 3), thick black and grey lines correspond to 1*l_x_* and 2*l_x_* domain sizes (see Appendix B for details on *l_x_*). Triangles represent highest (*S*_0_ = 5.7, 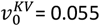) velocity gradients and squares represent lowest velocity gradients (*S*_0_ = 5.7, 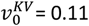) of the enhanced drag region presented in Figure 7.

The fact that the data from embryo experiments is fairly comparable to the curves resulting from our simulations suggest that our simulations are able to capture a range of velocity gradients that are similar to those found *in vivo*. Clearly, our experimental data is insufficient to pinpoint a specific rheology, although it is consistent with shear thinning behavior and obviously distinct from shear thickening behavior.

The similarities between normalized experimental and numerical velocity gradients indicate that the *rheologies* of the materials are similar. In contrast, the overall magnitudes of forces – such as those generated by shear stresses and pressure – are proportional to the large-scale effective viscosity of the tailbud, which is a parameter in the simulations that is unknown from experiments. If this viscosity is sufficiently large in experiments, our results predict that drag forces resulting from the observed PIV profiles would generate precisely the cell shape changes observed in experiments.

## 4. Discussion and conclusions

In this work, we develop a technically challenging 3D self-propelled Voronoi model with heterotypic surface tension to simulate confluent tissues. With this simulation code, we are able to use a physics-based model for individual cell shapes to efficiently simulate and predict the dynamic behavior of large groups of cells in 3D, including the temporal evolution of organ-scale structures. To our knowledge, this is the first time confluent tissue modeling on this scale has been attempted in three dimensions.

We model the motion of the zebrafish KV through the tailbud tissue for a range of tailbud tissue fluidities (*S*_0_) and KV initial velocities 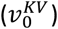 to understand how those parameters influence the AP distribution of KV cells (*n_a_/n_p_*). We find two regions in the parameter space in which the KV remains symmetric: (1) when the tailbud tissue is solid-like, the KV self-propulsion force is not strong enough to overcome forces imposed by the solid-like tailbud cells, and the KV does not move 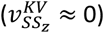; (2) when external cells are fluid-like, but the KV moves at such low speeds that forces on the KV remain very small. In the remaining parameter space (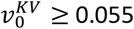 and *S*_0_ > 5.2) we found KV cell distribution values comparable to those found in experiments [11]: *n_a_/n_p_* = 1.77. The parameter scan allows us to focus the analysis on the region with enhanced asymmetry at the fluid-solid transition (*S*_0_ = 5.4 and 5.5) as well as an ostensibly fluid-like (*S*_0_ = 5.7) at various 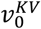.

We characterize streamlines of external cells around the spherical KV, along with pressure and shear stress exerted on the KV by external cells. Comparing our simulation results to the analytical solution of Stokes flow around a sphere, we identify dynamics that are similar to Newtonian flows as well as features indicative of non-Newtonian regimes. In the fluid-like regime (*S*_0_ = 5.5 and 5.7), streamlines, shear, and pressure behave similarly to Stokes flow, except at high 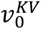. In the solid-like regime (*S*_0_ = 5.4), shear stresses and pressures are very different from Stokes predictions, indicating that more complex interactions occur.

Particle image velocimetry analysis of zebrafish embryos revealed a very robust velocity gradient around the KV, meaning that KV shape changes could be driven by mechanical forces generated by tissue velocity gradients around the KV. An important question is whether the natural variation in embryo-to-embryo velocity gradients would still allow robust shape changes. Although our experimental data do not directly answer this question, simulation results suggest the answer is yes. Specifically, Figure 5F shows that there is a very broad range of self-propulsion forces on the KV 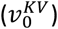 and tailbud cell shapes (*S*_0_) that all give rise to a *n_a_/n_p_* ratio between 1.4 and 2, which is what is observed in experiments. In other words, quite a large range of velocity gradients give rise to shape changes that are consistent with those required for downstream LR pattern formation.

Stronger velocity gradients are observed on the left side in the 2-4 ss and 4-6 ss compared 6-8 ss, suggesting the forces are stronger at earlier stages. Furthermore, our data (Figure 10E) hinted that there may be a decrease in drag forces from tailbud tissue to the left of the KV compared to the right, though more experiments will be needed to gain statistical confidence in this result. If this preliminary observation is confirmed, it is very exciting, as it suggests that LR asymmetry may exist at the tissue level upstream of all previously identified LR signals, including KV cell shape changes and directional flow inside the lumen.

Obviously, future work should focus on this possibility. It could be that the tailbud tissue of the left side is more viscous than the right side, possibly due to differing expression levels of myosin II. This is intriguing, because the higher viscosity would generate higher drag forces on the left side of the KV, which could potentially be sensed by mechano-sensory cilia. This process could work in parallel with the well-explained lumen directional flow that also triggers mechano-sensory cilia inside the KV to ensure that LR asymmetry is established. One possible set of future experiments to probe this hypothesis would be to optically excite caged Rho kinase inhibitors [44] to dynamically alter myosin II activity in a region of interest. This would allow one to alter the effective viscosity of tailbud cells on the left and right sides of the KV and determine whether such perturbations affect LR patterning. Such changes could also be implemented in simulations as a change to *S*_0_.

Another important conclusion of our work is that the KV motion acts as a probe of the tailbud tissue rheology. In simulations, we analyze the profiles of external cells velocity relative to the KV for various values of tailbud tissue fluidity *S*_0_. We find that the simulation velocity profiles are not Newtonian and instead behave similarly to viscoelastic flow around a sphere. An analysis of shear stress vs. rate of shear strain indicates that the external cells behave as shear-thinning viscoelastic fluid. We demonstrate that embryos’ normalized velocity profiles are comparable to those in simulations and consistent with shear thinning rheology. This means that the mechanical forces observed in our simulations are also present in the zebrafish embryo and would generate the observed KV cell shape changes if the effective viscosity of tailbud tissue is sufficiently large. In addition to studying the effect of tailbud tissue fluidity (*S*_0_) on the global rheology, a possible future direction is to investigate how the nonlinear global rheology of the tissue depends on vertex models parameters similarly to the works of Tong, Singh [45], Duclut, Paijmans [46].

One obvious implication of this work is that a direct measurement of the effective viscosity of zebrafish tailbud tissue, in response to large-scale perturbations, would be particularly valuable. The challenge is that existing methods [11, 12] focus on relatively small-scale (i.e. cell scale) perturbations to embedded droplets, which means that they do not probe the tissue-scale effective viscosity generated by many cell neighbor exchanges in the presence of large flows or perturbations. Previous work on materials such as foams, emulsions [23], and even mucus [47], suggest that these microscopic and macroscopic rheology measurements often generate very different results.

Another important question is whether our results are sensitive to the type of model we simulated. In this manuscript, we focused on a confluent model for tailbud tissue, although recent work has demonstrated that there are small gaps between cells in the tailbud, and even suggested that particle-based models for cell packings may better capture the nature of the fluid-solid transition in that region [15]. Importantly, the drag forces that we find here are likely to be quite generic across models, as both Voronoi and particle-based foam models [48] are shear thinning and are close to a fluid-solid transition in the parameter regime relevant for tailbud tissue. Therefore, flows in any of these models – consistent with the experimentally observed streamlines – will generate similar drag forces up to an overall scaling factor related to the effective viscosity. Moreover, the epithelial nature of KV cells suggests that a confluent model will best capture how drag forces affect cell shapes within the KV.

Another related consideration is that the model presented here possesses highly simplified shapes – the lumen and KV layer were initially spherical, and we do not represent other structures such as the notochord, yolk, or tailbud boundary. We made this choice to focus on how the rheology of the full 3D Voronoi model (which represents each individual cell) generates drag. However, now that we have demonstrated the Voronoi model rheology is relatively well-captured by relatively simple viscoelastic constitutive laws, this opens the door to a different type of simulation that can account for the complicated geometry of the tailbud. Future work could focus on a computational fluid dynamics (CFD) approach, where the external cells are modeled as a continuum fluid and the KV is modeled as a roughly spherical deformable elastic object. In this context, we could investigate how different rheology models for the tailbud tissue, as well as the presence of boundaries generated by the yolk and tailbud boundary, affect the overall shape of the KV and the AP asymmetry of cell shapes.

Finally, this work has significant implications beyond the simple KV organ we study here. The computational tools we have developed can be immediately applied to predict the dynamics of other 3D structures during development and disease processes with cellular resolution. In particular, we have demonstrated that PIV analysis of experimental data to extract streamlines combined with computational modeling can help provide a strong indication of when dynamic, fluid-like forces are acting to shape organs during development. To our knowledge, such dynamic forces have been rarely studied [13, 20, 49], but they are likely to be important in other processes where groups of cells migrate across a tissue.

## Supporting information

Revised Supplementary Information

Revised Supplementary Video S1

Revised Supplementary Video S3

Revised Supplementary Video S4

## Appendix A. Additional details for PIV analysis of experimental data

In our Particle Image Velocimetry processing, each image is divided into subregions (interrogation areas) and correlation functions determine the displacement of subregions between consecutive frames. To reduce errors, we iterate this procedure with decreasing interrogation area [50]. Displacements are then converted to velocity fields using the known frame rate (frame/minute) and pixel-to-micron ratio. Due to the different experimental setups between embryos, e.g. varying frame rates and objectives, the size of interrogation areas and number passes were optimized for each embryo. For all embryos, the smallest interrogation area is about the size of a tailbud cell nucleus (e.g. 32 pixels for 20X objective). After velocity fields (***v**_PIV_*) are obtained, very few data points are deemed outliers (*v_PIV_x__, v_PIV_z__* ≥ 10 *μ*m/min), which are interpolated using the “vector validation” tool of PIVlab [28].

We determine KV velocity, in each frame, from the PIV analysis 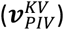 by averaging velocity vectors (***v**_PIV_*) within the KV area. The KV area is manually determined in each frame using ImageJ’s [25] tools “oval” and “ROI Manager”. Then, tailbud cells velocity relative to KV is calculated for each frame: 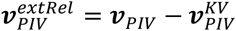.

In order to average velocity fields over multiple frames, we use the KV at the mid-frame (center frame of *n_frames_*) as the KV reference frame and subtracted KV centroid from the position of velocity vectors in all frames. The centroid is obtained by using the center of the KV areas as described above. Then, velocity fields of all frames are averaged over the complete 50-minute time window (Figure 3) – we used *n_frames_* that corresponds ~50 min (or 2 ss in our laboratory conditions). With each embryo’s averaged velocity field 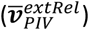 on a regular grid, streamlines for 2 ss are calculated using MATLAB’s [51] function “streamslice.” Velocity profiles are obtained by interpolating the averaged velocity field 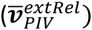 at points along the left-right axis (deemed *x*-axis) that passes through the KV centroid and analyzing the *z*-component of velocity 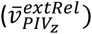 along the *x*-axis.

## Appendix B. Additional details for the 3D SPV simulations

As the fluid-filled lumen is expected to be incompressible, the lumen volume elasticity (*K_V_*) is set higher than those of other cell types, i.e. lumen cells were less compressible than other cell types and their volume remained fairly constant during simulations. Although the lumen expands during embryonic development, volume expansion has been experimentally demonstrated to be irrelevant for changes in shape of KV cells [6] and so we avoid that complication in our simulations. To avoid rearrangements between lumen and KV cells, the preferred shape index of KV cells is set to be in the solid regime. A surface tension, *γ*, was added along the interface between different cell types: *γ*_KV-external_ =1 *τ*_KV-lumen_=1, *γ*_lumen-external_=100. We choose to place a very high interfacial tension between lumen and external cells (a) to ensure that they do not mix and (b) to retain the lumen spherical shape as observed in experiments. The preferred surface area of external cells (*S*_0_) varies from 5.0 to 6.0 (Figure 4D) to represent the various tissue fluidities. Table B1 summarizes the parameters used in the simulations.

**Table B1:** Simulation parameters in natural units

| | N | Volume elasticity $K_V$ (-) | Preferred volume $V_0$ ($l^3$) | Surface elasticity $K_S$ (-) | Preferred surface $S_0$ ($l^2$) |
| --- | --- | --- | --- | --- | --- |
| External | 2048 | 1 | 1 | 1 | 5.0-6.0 |
| Lumen | 17 | 10 | 1 | 1 | 5.0 |
| KV anterior | 25 | 1 | 1 | 1 | 5.0 |
| KV posterior | 25 | 1 | 1 | 1 | 5.0 |

Each simulation consists of two stages: First, Equation (2) is integrated in the absence of any self-propulsion (*v*^0^ = 0) using Euler’s method (*dt* = 0.01*τ*) until the system converges to a state of minimal energy (we define convergence by defining the maximum energy of an individual cell in an iteration and then calculating the average maximum cell energy for five consecutive iterations; convergence is achieved when the difference between averages is less than 10^-5^). Second, a fixed self-propulsion speed 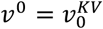, along a fixed direction 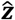 is introduced to every cell in the KV (i.e. KV and lumen cells), and Equation (*2*) is integrated using Euler’s method (*dt* = 0.001*τ*) for all cells in the simulation. The self-propulsion forces on the KV causes the entire KV to move through the surrounding tissue, and we measure the resulting speed of the KV centroid (*v^KV^*), which is slower than 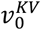 due to interactions with the tailbud. We define the simulation as being in steady state (SS) when the observed KV instantaneous speed 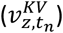 has converged, so that differences in observed speed between simulation time points drop below a threshold of 5×10^-7^. KV instantaneous speed is the difference between the KV centroid *z*-position 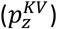 at two consecutive time steps 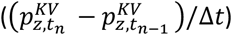. We observed that when the external cells are clearly solid-like (*S*_0_ < 5.4), SS occurs after about 100,000 iterations and in the fluid-to-solid transition as well as fluid-like regime, SS occurs after about 200,000 iterations (Supplementary Figure S8 of steady-state velocity convergence). To ensure we record significant data beyond the start of steady state dynamics, we run simulations for 200,000 iterations when *S*_0_ < 5.4 and 400,000 iterations when *S*_0_ < 5.4.

In zebrafish embryos, the tailbud tissue does not move along with the KV, most likely because the tailbud tissue at the boundary of the tailbud does not move relative to the edge of the embryo, known as a no-slip boundary condition in fluid dynamics. It is also possible that more complicated pinning at the yolk interface or with other structures occurs. To mimic this effect in our simulations and prevent the entire system from translating in the direction of the self-propulsion velocity 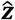, 2% external cells farthest from the center of the KV and parallel to 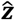 were pinned by fixing their position at every time step (only when *v*_0_ is added). In the fluid-like regime, we have checked that the precise density (1, 2, or 4%) of pinned cells does not significantly impact the results in the fluid-like regime. In the small region of phase space where the tissue response is viscoelastic (discussed in the results Section 3.1), the pinning density does alter the quantitative results, although the qualitative response remains similar independently of pinning density. This is not surprising, given long-range elastic interactions in viscoelastic materials, and is an interesting avenue for future work. For simplicity we fix the pinning density at 2% in viscoelastic simulations, as that is sufficient to prevent the tailbud from translating along with the KV.

Simulations of 3D cells are substantially more computationally expensive than 2D. Thus, to avoid unnecessary use of computing resources, we determine the smallest domain that would allow simulations where the periodic copies would not affect the macroscopic KV motion. We test three cubic domain sizes with increasing number of external cells (*N_ext_* = 1028, 2048, and 4096), compared KV traveled distance, and concluded that *N_ext_* = 2048 is sufficiently large such that the box size does not interfere in the KV traveled distance. We denote this domain the “regular” domain size with side of length *l_x_*. In a few instances, we also simulate larger system sizes (by increasing *N_ext_* such that the *x* and *y* dimensions doubles, but *z* dimension remains the same), which we denote the larger domain size as 2*l_x_*. Data presented in this paper reflect the average of ~30 simulations with different initial conditions for each pair of parameters (*S*_0_ and 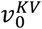). In very few instances, simulations were discarded when a Voronoi tessellation could not be determined by the voro++ library [52] due to a degenerate higher-fold coordinated vertex that is incompatible with our data structure, which generates a run-time error.

## Appendix C. Streamline calculations in the Voronoi model simulations

In order to obtain streamlines of external cells around KV (and eventually velocity gradients), several calculations are performed. First, we calculate the velocity of individual external cells *i* between simulation frames 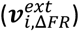 with a frame rate Δ*FR* of 5,000 iterations (here, frame rate refers to how often output files are generated). Second, we compute the KV centroid as the average position of lumen cells, and its velocity between simulation frames 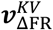. Third, we obtain the velocity of individual external cells relative to the KV velocity (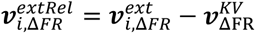), which generates an unstructured velocity field. Fourth, to average velocity fields of multiple simulations, we create a structured spherical grid with length scale normalized by KV radius (*r_KV_*), i.e., KV is represented as a unit sphere. Figure C1B shows the spherical structured grid with polar angle: 0 < *θ* < *π* and Δ*θ* = *π*/12; azimuth angle: 0 < ϕ < 2*π* and Δ*ϕ* = *π*/24; and radial axis: 1 ≤ *r* < 3.5 *r_KV_* and Δ*r* ≈ 0.25 with spherical coordinates defined in Figure C1A. We obtain the KV radius by calculating the KV total volume and approximating it as a sphere to find its radius. Fifth, the unstructured velocity field of external cells 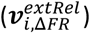 is interpolated to the structured spherical grid. At this stage, we have (1) fixed the KV as a unit sphere with center at the origin and (2) placed external cells velocity relative to the KV on a structured spherical grid. Sixth, for each simulation, we calculate the average velocity field for 10 frames after steady state is reached. Finally, we average ~30 simulations per ensemble to calculate the averaged structured velocity field of external cells per ensemble (Figure C1D).

**Figure C1:**
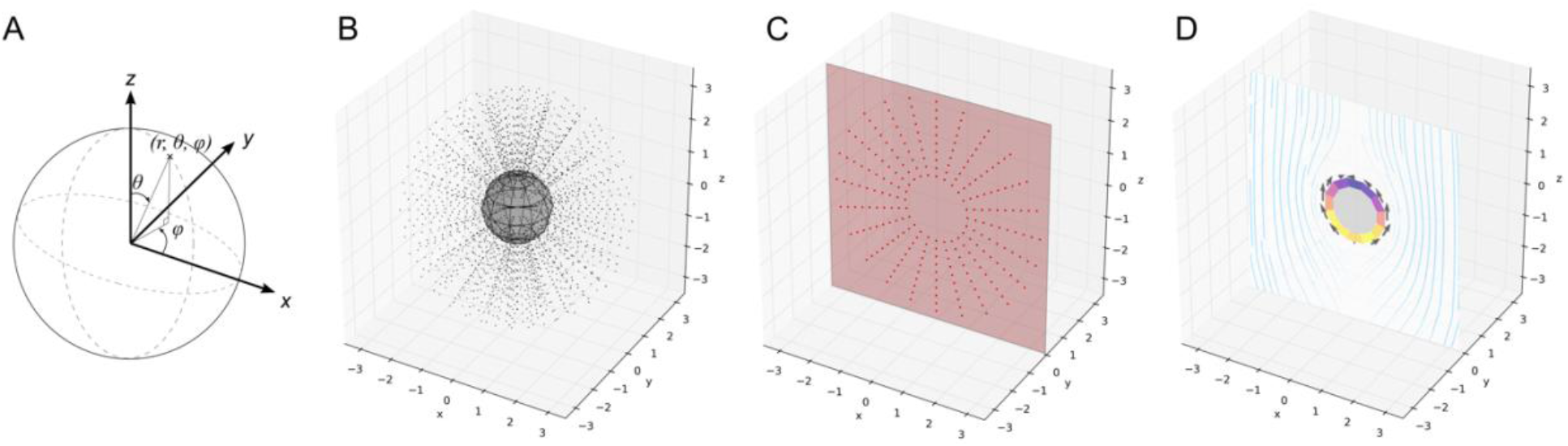
Structured spherical grid used to generate 2D streamlines for ease of analysis. (A) Spherical coordinate system with radial distance *r*, polar angle *θ* (latitude), and azimuth angle *ϕ* (longitude). (B) Black dots are the points on the 3D spherical structured grid with KV represented as a unit sphere. (C) Red dots represent points that are rotated around the azimuth angle *ϕ* that lay on the red 2D plane (*xz* plane, *y* = 0). (D) Calculated streamlines, pressure and shear on the 2D plane.

In this simulation geometry the flow is axisymmetric [14, 53, 54], and so we simplify our analysis to 2D by rotating velocity vectors along the azimuth (*ϕ*) to the *xz*-plane and averaging vectors that belong to the same position (Figure C1C). Then, we calculate the projection of 3D velocity vectors onto the *xz*-plane to obtain 2D vectors. At this final stage, we have averaged 10 frames per simulation; ~30 simulations per ensemble; and 12 angles around the azimuth. For simplicity, we denote 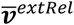 this final average of external cells velocity relative to the KV on the structured grid. Finally, streamlines shown in Figure C1D are calculated using Matplotlib [55] function streamplot. The procedures just described for transforming 3D unstructured velocity field onto the final 2D plane of Figure C1D were also performed for the pressure and shear, which are represented by the circular rim and arrows, respectively. Velocity profiles are obtained by using the *z*-component of external cell velocity 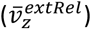 at the KV equator line (*θ* = *π*/2) and at KV poles (*θ* = 0,*π*). In order to compare simulations of different parameters, 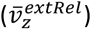 is normalized by the initial self-propulsion velocity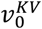. To compare velocity profiles of simulations and experimental results, we use the embryo tailbud cell diameter of ~10 *μm* to convert the average size of an external cell in the simulation in natural units.

## Acknowledgements

We would like to thank Matthias Merkel for his technical assistance as well as developing and sharing the original version of the 3D Voronoi code.

## Funding

This work was supported by NIH grants R01GM117598 and R01HD099031, and a Simons Foundation Investigator Award (#446222).

## Notes

### Competing Interest Statement

The authors have declared no competing interest.

### Summary of Updates

This version of the manuscript has been revised to update the following: embryo movies, PIV analyses, and PIV analysis error estimate.

