## Supplementary material for "3D viscoelastic drag forces contribute to cell shape changes during organogenesis in the zebrafish embryo": Revised Supplementary Information

### 1. Supplementary tables

Table S1: Experimental details of each embryo

| Embryo | Objective | Z-step (um) | Time interval (min) | Starting stage | Adjusted starting time (min) | Number of frames | Duration (min) | Adjusted ending time (min) | Ending stage |
|---|---|---|---|---|---|---|---|---|---|
| Embryo 01 | 20X | 2 | 2 | 4 ss | 50 | 65 | 128 | 178 | 9 ss |
| Embryo 02 | 20X | 2 | 4 | 3 ss | 25 | 41 | 160 | 185 | 9 ss |
| Embryo 03 | 20X | 5 | 2 | 4 ss | 50 | 31 | 60 | 110 | 6 ss |
| Embryo 04 | 20X | 3 | 2 | 4 ss | 50 | 51 | 100 | 150 | 8 ss |
| Embryo 05 | 20X | 3 | 2 | 2 ss | 0 | 57 | 110 | 110 | 6 ss |
| Embryo 06 | 20X | 3 | 2 | 3 ss | 25 | 52 | 100 | 125 | 7 ss |
| Embryo 07 | 20X | 3 | 2 | 2 ss | 0 | 43 | 84 | 84 | 5 ss |
| Embryo 08 | 20X | 2 | 2 | 2 ss | 0 | 60 | 118 | 118 | 6 ss |
| Embryo 09 | 20X | 2 | 2 | 3 ss | 25 | 56 | 110 | 135 | 7 ss |
| Embryo 10 | 20X | 2 | 2 | 4 ss | 50 | 44 | 86 | 136 | 7 ss |
| Embryo 11 | 20X | 2 | 2 | 2 ss | 0 | 26 | 50 | 50 | 4 ss |
| Embryo 12 | 20X | 2 | 2 | 6 ss | 100 | 34 | 66 | 166 | 8 ss |
| Embryo 13 | 20X | 2 | 2 | 2 ss | 0 | 48 | 94 | 94 | 5 ss |
| Embryo 14 | 20X | 2 | 2 | 6 ss | 100 | 30 | 58 | 158 | 8 ss |

Table S2: KV features variation reported as one standard deviation as a percentage of the mean: left and right fitted velocity gradient of individual embryos and KV size (maximum cross-sectional area of lumen) and cilia number. *values obtained from Gokey, Ji [1] correspond to the end of the development window (e.g. value for 2-4ss was measured at the 4 ss).

| | Fitted gradient | | | |
|---|---|---|---|---|
| Somite stage | Left | -Right | KV size* | Cilia number* |
| 2-4 | 27.1% | 32.9% | 44.10% | 33.30% |
| 4-6 | 71.5% | 54.8% | 41.40% | 42.30% |
| 6-8 | 46.5% | 69.4% | 34.00% | 30.60% |

#### 2. Supplementary figures

**KV and temperature methods:** To test the impact of incubating *Tg(sox17:GFP-CAAX)* embryos at 32°C during live imaging, embryos were either maintained at 28.5°C (control) or shifted from 28.5 to 32°C from 2 ss to 8 ss (temp shift). Embryos were then mounted at 8 ss to image KV or returned to 28.5°C until 2 days post-fertilization to score heart looping. To quantify KV size, we measured $KV^{max}$ area, given by the area of the KV lumen at its maximum diameter, using ImageJ software (NIH) as previously described [1]. Heart looping was scored in live embryos as normal, reversed, or no looping.

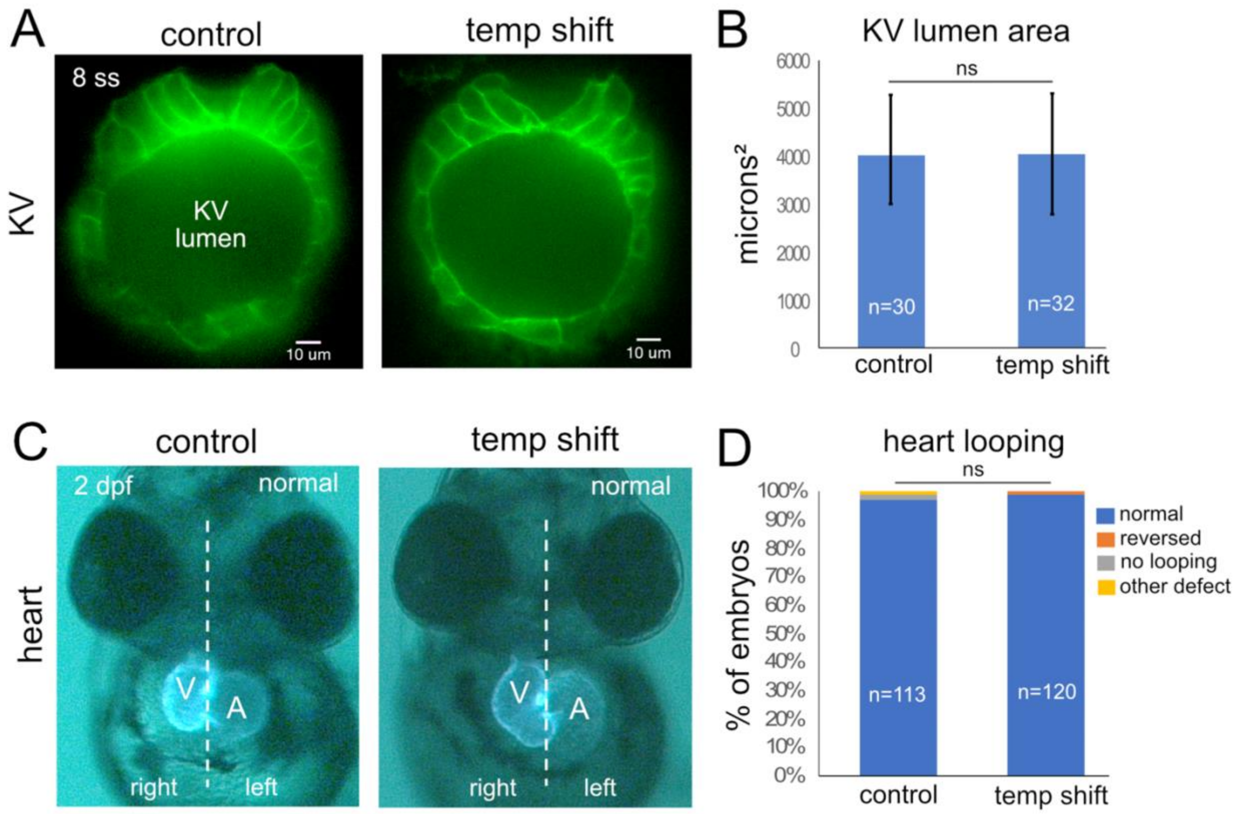


Figure S1: A temperature shift to 32°C does not alter KV development or left-right patterning in *Tg(sox17:GFP-CAAX)* embryos. (A) Representative images of the middle plain of KV at 8 ss in *Tg(sox17:GFP-CAAX)* embryos that were either maintained at 28.5°C (control) or shifted from 28.5 to 32°C from 2 ss to 8 ss (temp shift). (B) Quantification of $KV^{max}$ area (area of the lumen at its maximum diameter) at 8 ss in control and temp shift embryos pooled from three independent experiments. (C) Normal control and temp shift embryos (shifted from 28.5 to 32°C from 2 ss to 8 ss and then returned to 28.5°C) at 2 days post-fertilization (2 dpf). During normal heart looping, the ventricle (V) is positioned on the right and the atrium (A) is on the left. Cardiomyocytes in the heart are labeled with GFP by *Tg(myl7:GFP)* expression in *Tg(sox17:GFP-CAAX)* embryos. Dashed line is embryo midline. (D) Heart looping in control and temp shift embryos. n=number of embryos analyzed. ns=no significant difference (unpaired t test with Welch's correction). Error bars=1 standard deviation.

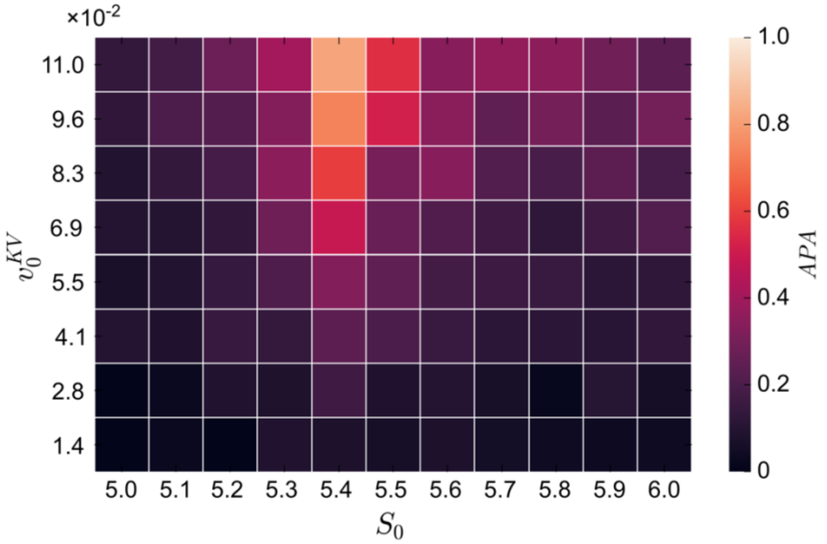


Figure S2: Sweep in parameter space of tailbud tissue fluidity ($S_0$) and KV initial velocity ($v_0^{KV}$ = 0.01375, 0.0275, 0.04125, 0.055, 0.06875, 0.0825, 0.09625, and 0.11 labeled for simplicity as 1.4, 2.8, 4.1, 5.5, 6.9, 8.3, 9.6, 11 × 10$^{-2}$) for $APA = LWR_{ant} - LWR_{post}$.

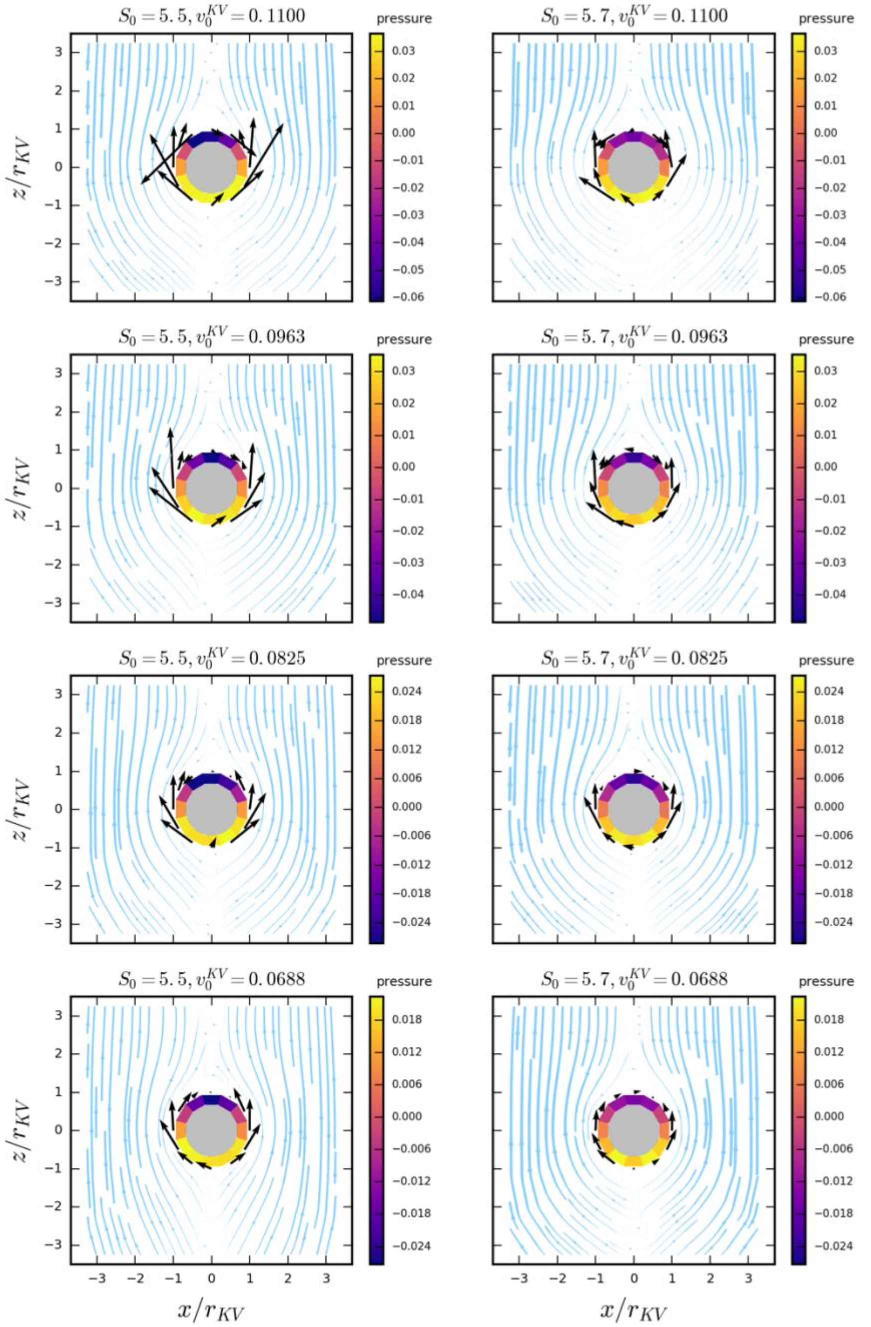


Figure S3: Streamlines, pressure, and shear for various tailbud tissue fluidities $S_0$ and initial KV velocities $v_0^{KV}$. (figure continues on next page).

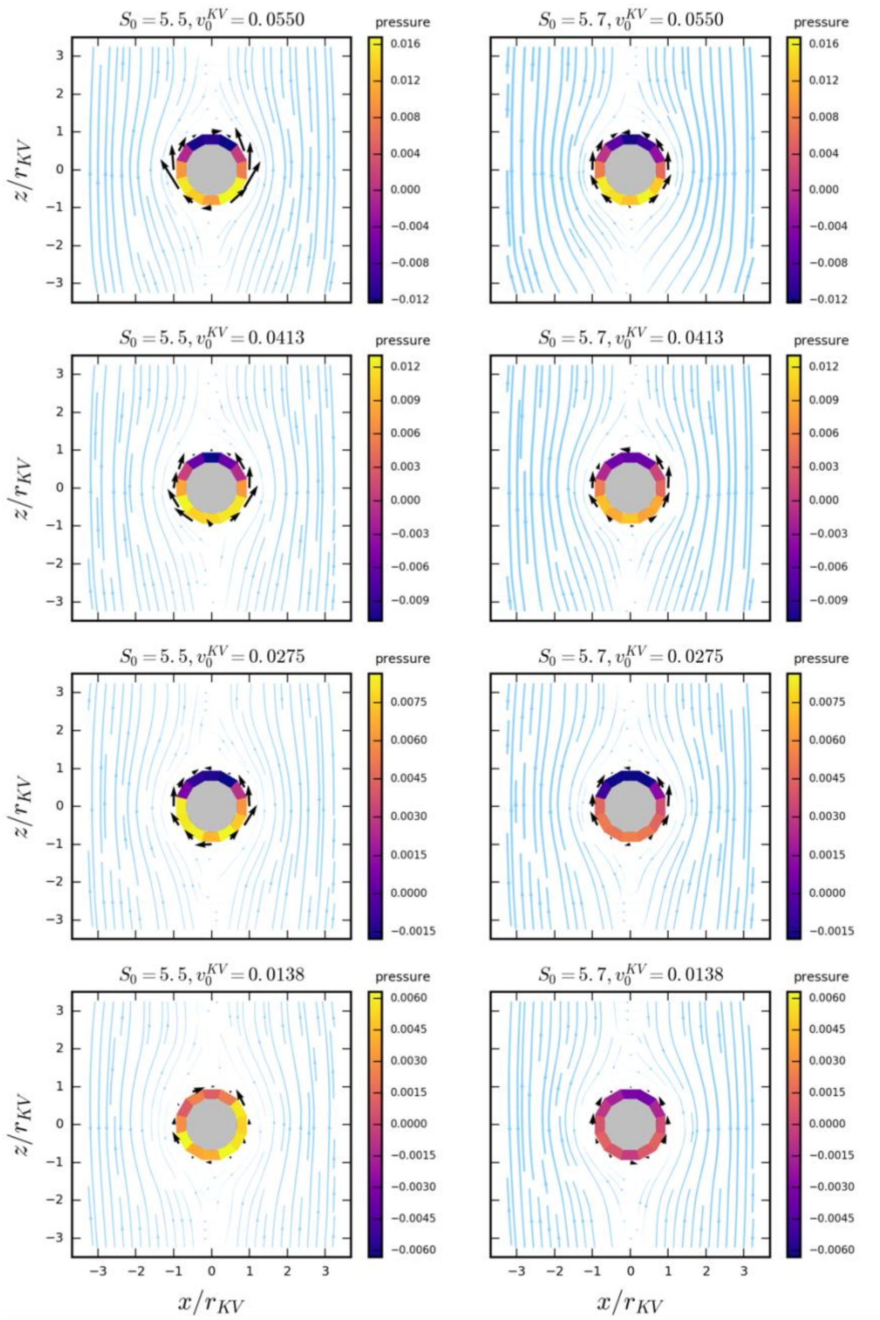


Figure S3: (continued from previous page) Streamlines, pressure, and shear ensemble average for various tailbud tissue fluidities $S_0$ and initial KV velocities $v_0^{KV}$. Each row is the result of a fixed $v_0^{KV}$ value whereas each column is the result for a given $S_0$. The colored ring represents the pressure exerted by the external cell fluid flow on the sphere, with positive pressure (pushing inward, yellow- ranged colors) negative pressure (pulling outward, purple-ranged colors). Note that the range of the pressure colorbar is fixed per row and, thus, pressure colors can be compared withing a row. Arrows represent the shear stress acting tangentially on the KV (the arrow length represents the magnitude with the same scale for all plots, and the location of the shear stress is at the tail of the arrow). $x$ and $z$ axes are normalized the KV radius, $r_{KV}$ (for more details on this normalization, refer to Appendix D of manuscript). Streamline width represents the velocity of external cells relative to the KV ($\bar{v}^{extRel}$) normalized by the initial KV velocity ($v_0^{KV}$) such that streamline widths are comparable within a row.

**PIV error estimate:** Although we cannot calculate PIV errors by comparing the PIV results of *in vivo* data to a specific standard such as analytical solution or measurements of velocity from probes in the embryo, we can roughly estimate systematic errors (e.g. in regions where there is little image signal the cross-correlation could identify a flow that is an artifact instead of real motion of cells). Specifically, we calculated local fluctuations of the velocity magnitude. Considering the laminar flow in the tailbud, we expect only small differences between velocity magnitudes in nearby points on the grid. Therefore, large local fluctuations are an indication that the PIV analysis is picking up noise rather than signal. For this reason, the magnitude of error in the extracted velocity fluctuations provides a reasonable estimate of this systematic error in the PIV analysis. Similar to the calculation performed in [2], we calculated the local average by averaging the velocity magnitude within a radius of 1.5x of the smallest PIV interrogation area (for most cases, the smallest interrogation area was 32 pixels). In Figure S4 we present the local average of Embryo 10's velocity field (Figure 9A of manuscript) field and its corresponding error. The error at the smallest interrogation area $i, j$ is calculated by the absolute value of the difference between local average velocity magnitude and the PIV velocity magnitude: $PIV_{error} = \left| \left\| v_{localAvg_{i,j}}^{extRel} \right\| - \left\| v_{PIV_{i,j}}^{extRel} \right\| \right|$.

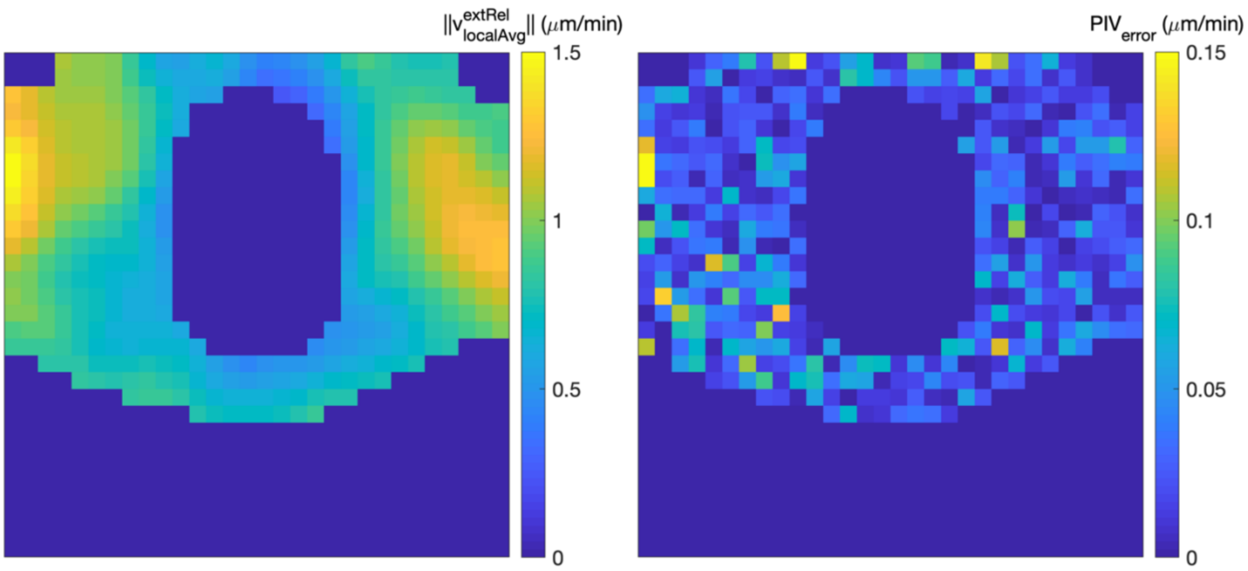


Figure S4: Estimation of PIV error from the PIV analysis of Embryo 10 (Figure 9A of manuscript). Left: velocity magnitude local average $v_{localAvg_{i,j}}^{extRel}$ for the smallest interrogation area $i, j$. Right: PIV error estimate, $PIV_{error}$.

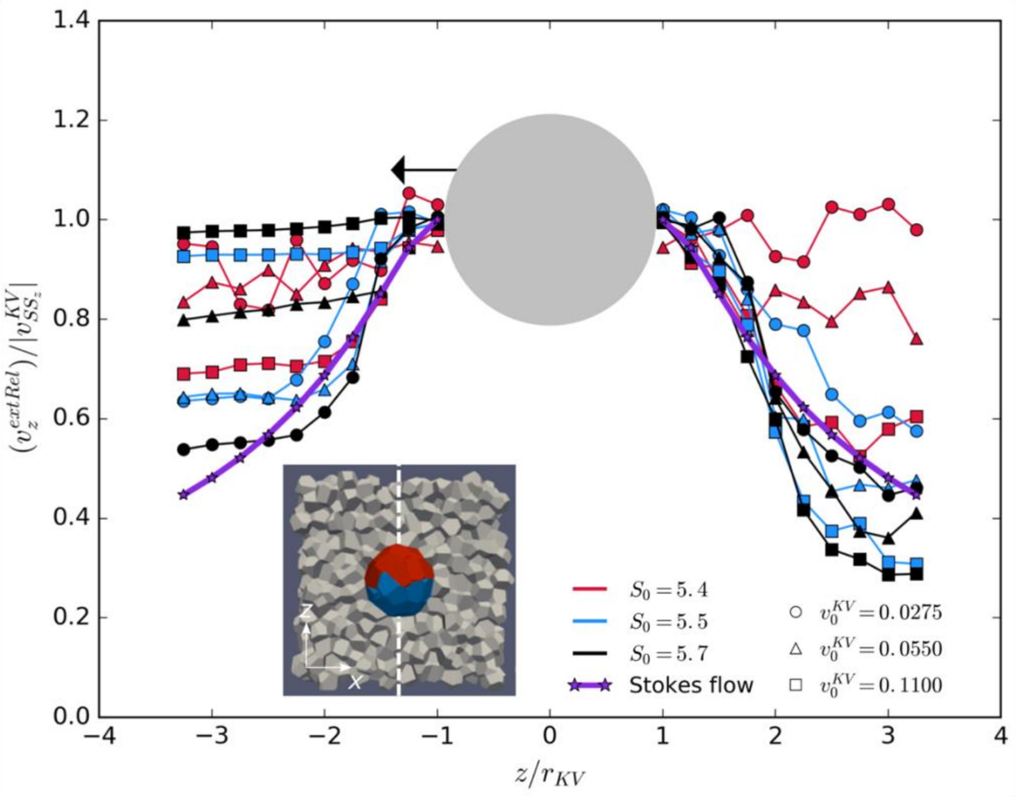


Figure S5: Wake profile, $v_z$, for the regular domains of ~$3.5r_{KV} = 1l_x$ at the poles as indicated in the inset.

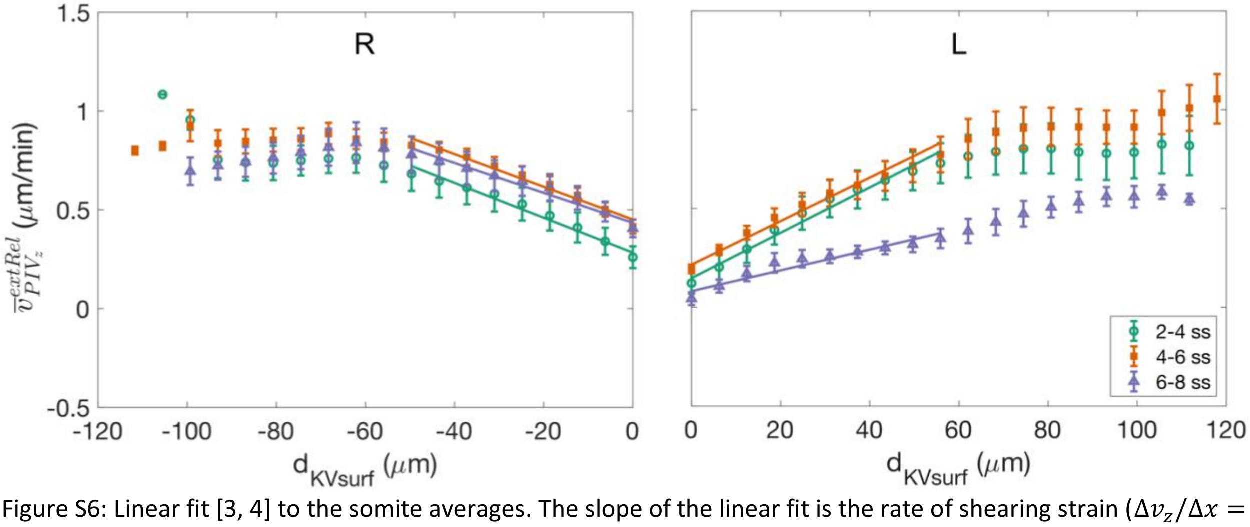


Figure S6: Linear fit [3, 4] to the somite averages. The slope of the linear fit is the rate of shearing strain ($\Delta v_z/\Delta x = \dot{\gamma}$ s$^{-1}$) and should be proportional to the drag forces exerted by tailbud tissue on the KV. The fit only includes points within ~50 $\mu$m from the KV surface due to our interest in the drag forces near the KV. $v_{PIV_z}^{extRel}$ is the external cells $z$-component of velocity relative to the KV and $d_{KVsurf}$ is the distance from the KV surface.

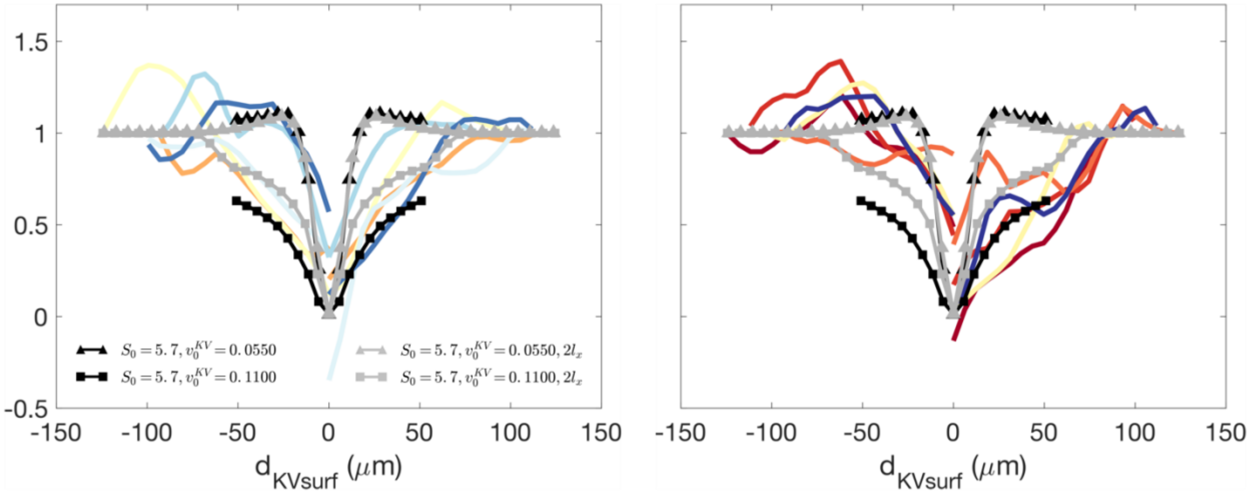


Figure S7: Normalized velocity profiles with embryos represented by colored lines (each color represents embryos defined in Figure 3 of the manuscript), thick black and grey lines correspond to $1l_x$ and $2l_x$ domain sizes (see Appendix B for details on $l_x$). Triangles represent highest ($S_0 = 5.7$, $v_0^{KV}$= 0.055) velocity gradients and squares represent lowest velocity gradients ($S_0 = 5.7$, $v_0^{KV}$= 0.11) of the enhanced drag region presented in Figure 7 of the manuscript.

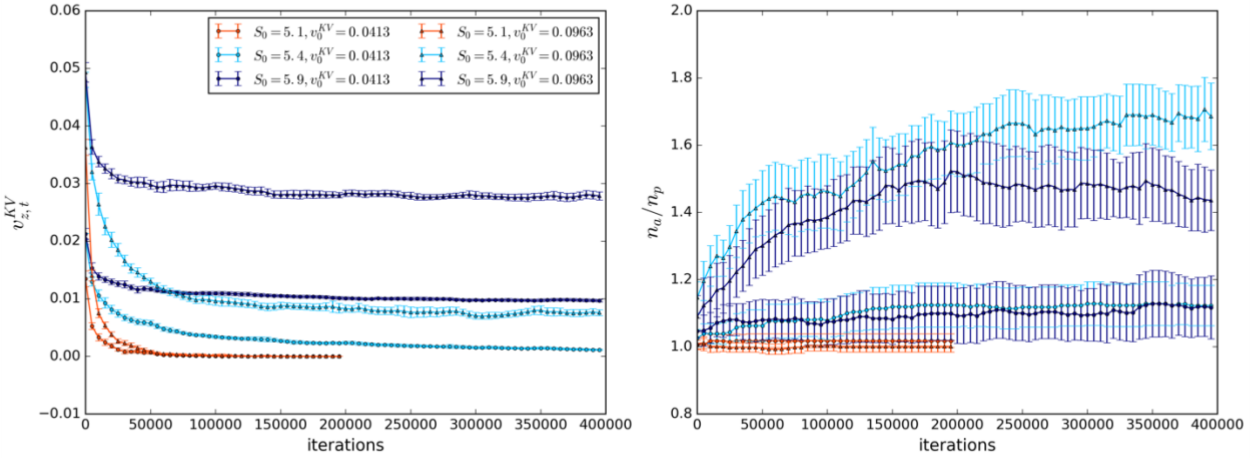


Figure S8: Ensemble averages at each iteration when KV self-propulsion velocity ($v_0^{KV}$) is included for different tailbud tissue fluidity ($S_0$) and KV self-propulsion velocities ($v_0^{KV}$). Left: $z$-component of KV instantaneous velocity ($v_{z,t_n}^{KV}$). Right: $n_a/n_p$ (right). The convergences of $v_{z,t_n}^{KV}$ and $n_a/n_p$ show that the systems reached state state.

#### 3. Supplementary videos

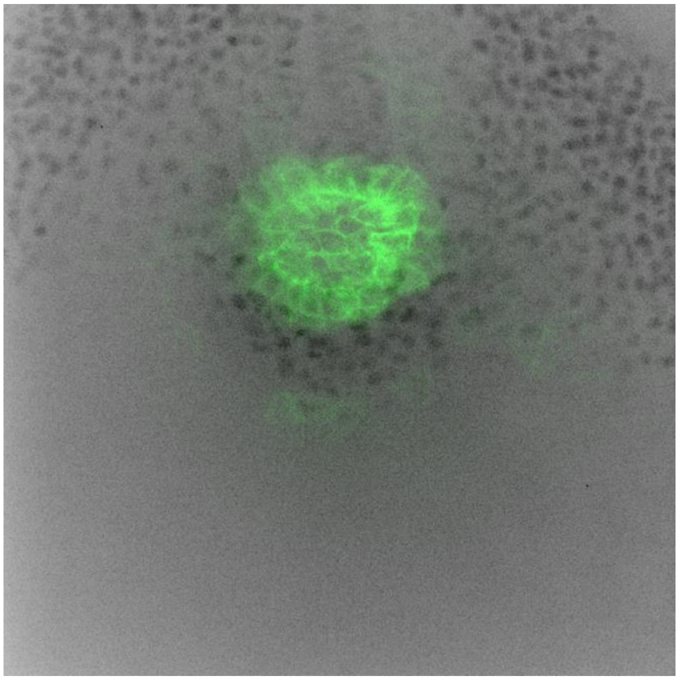

**Supplementary Video S1:** Live imaging between 4 and 7 ss of Embryo 10, a transgenic Tg(sox17:GFP-CAAX) zebrafish embryo with fluorescently labeled KV and external cell nuclei labeled by expressing nuclear-localized mCherry proteins.
File: SupplementaryVideoS1_Embryo10.avi.

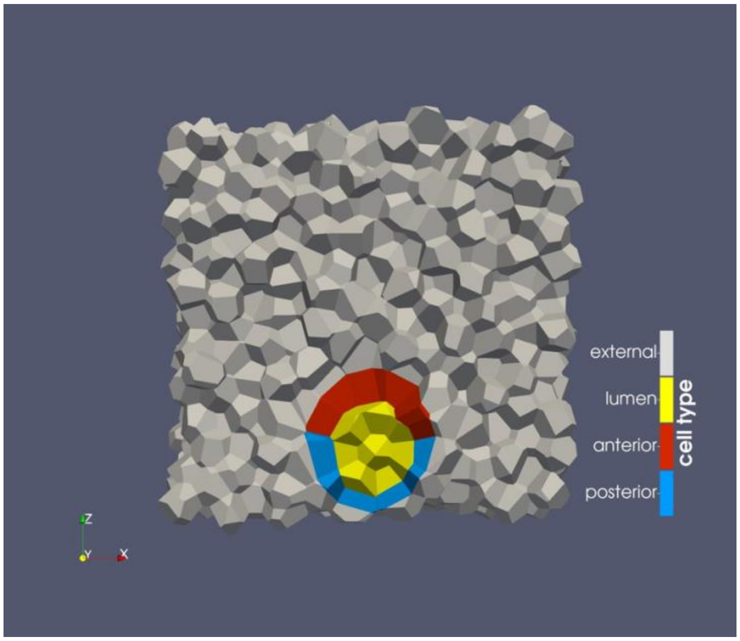


**Supplementary Video S2:** Simulation of KV moving through external cells that generates KV cell shape changes. Simulation parameters: $S_0$ = 5.5, $v_0^{KV}$ = 0.09625, $N_{ext}$ = 2048.
File: SupplementaryVideoS2_Simulation.avi.

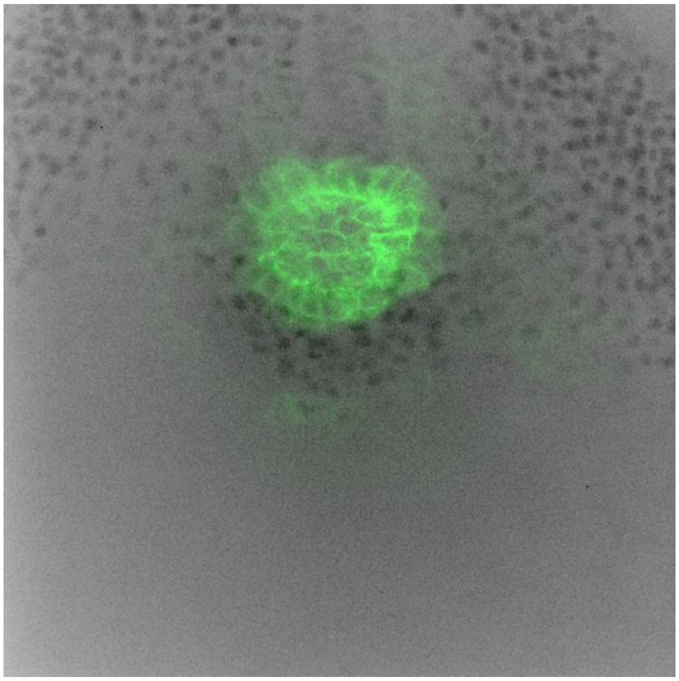

**Supplementary Video S3:** Fixing the KV as the reference object of Supplementary Video S1: each frame has been translated in space so that the KV position remains fixed as a reference object.
File: SupplementaryVideoS3_Embryo10_translated.avi.

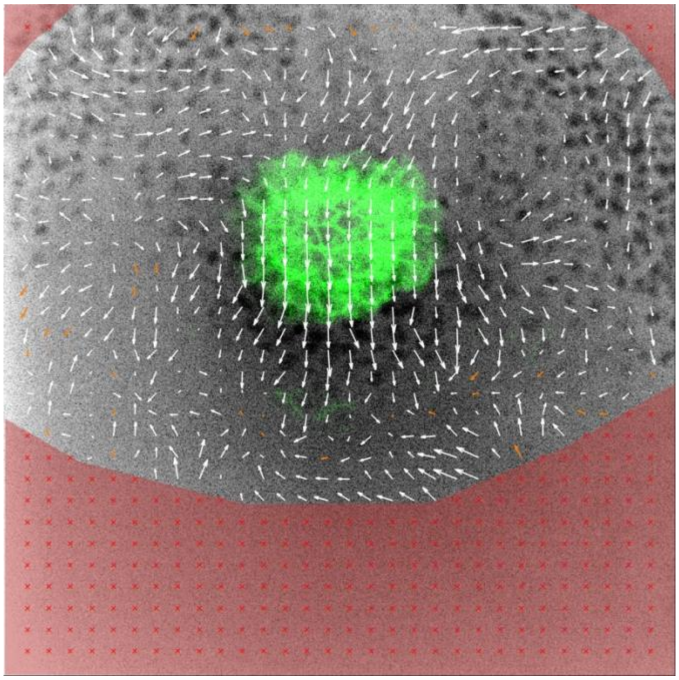

**Supplementary Video S4:** PIV analysis of Embryo 10 between 4 and 6 ss.
File: SupplementaryVideoS4_Embryo10_PIV_4-6ss.avi.
